# The type 1 diabetes-associated lncRNA *ARGI* participates in virus-induced pancreatic β cell inflammation

**DOI:** 10.1101/2022.12.01.518685

**Authors:** Itziar González-Moro, Koldo Garcia-Etxebarria, Luis Manuel Mendoza, Nora Fernández-Jiménez, Jon Mentxaka-Salgado, Ane Olazagoitia-Garmendia, María Nicol Arroyo, Toshiaki Sawatani, Anne Op de Beek, Miriam Cnop, Mariana Igoillo-Esteve, Izortze Santin

**Affiliations:** Department of Biochemistry and Molecular Biology, University of the Basque Country, 48940 Leioa, Spain; Biocruces Bizkaia Health Research Institute, 48903 Barakaldo, Spain; Biodonostia, Gastrointestinal Genetics Group, 20014 San Sebastián, Spain; Centro de Investigación Biomédica en Red de Enfermedades Hepáticas y Digestivas (CIBERehd), 08036 Barcelona, Spain; Department of Genetics, Physical Anthropology and Animal Physiology, University of the Basque Country, 48940 Leioa, Spain; ULB Center for Diabetes Research, Université Libre de Bruxelles, 1070 Brussels, Belgium; Division of Endocrinology, Erasmus Hospital, Université Libre de Bruxelles, 1070 Brussels, Belgium; Centro de Investigación Biomédica en Red de Diabetes y Enfermedades Metabólicas Asociadas (CIBERDEM), Instituto de Salud Carlos III, 28029 Madrid, Spain

**Keywords:** Interferon-stimulated genes, long non-coding RNA, pancreatic β cells, type 1 diabetes, viral infection

## Abstract

Type 1 diabetes-associated single nucleotide polymorphisms are mainly located in non-coding regions of the human genome. Single nucleotide polymorphisms located in long non-coding RNAs may result in the disruption of their secondary structure, affecting their function.

Here, we functionally characterized the virus-induced type 1 diabetes-associated lncRNA *ARGI* (Antiviral Response Gene Inducer). *ARGI* upregulation in pancreatic β cells leads to the transcriptional activation of antiviral and pro-inflammatory genes. Upon a viral insult, *ARGI* is upregulated in the nuclei of pancreatic β cells and binds to CTCF to interact with the regulatory regions of *IFNβ* and interferon-stimulated genes, promoting their transcriptional activation in an allele-specific manner.

The presence of the risk allele for type 1 diabetes in *ARGI* induces an hyperactivation of type I IFN response in β cells, an expression signature that is present in the pancreas of diabetic patients. These data shed light on the molecular mechanisms by which type 1 diabetes-related single nucleotide polymorphisms in long non-coding RNAs influence pathogenesis at the pancreatic β cell level.

## Introduction

Type 1 diabetes (T1D) is a chronic autoimmune disease characterized by the specific destruction of insulin-producing pancreatic β cells that leads to impaired insulin production and increased blood glucose levels^1^. During the initial stages of the disease, immune cells infiltrate pancreatic islets, generating a pro-inflammatory environment (insulitis) that is tightly controlled by the release of soluble pro-inflammatory mediators by both pancreatic β cells and immune cells^2^.

Over the past years, several studies have pointed to the role of T1D candidate genes in the regulation of the pro-inflammatory process that precedes the autoimmune destruction of pancreatic β cells in T1D^3-6^. The molecular mechanisms by which most T1D candidate genes influence T1D pathogenesis remain to be clarified. Accumulating evidence suggests that T1D risk genes interplay with viral infections in pancreatic β cells, promoting an imbalanced antiviral and pro-inflammatory response, that culminates in the autoimmune destruction of insulin-producing cells^3-6^. Indeed, the role of viral infections in T1D development is supported by clinical and epidemiological data^7-12^.

Although inflammation- and virus-induced activation of specific transcription factors (IRF7, NFκB, STAT1 and STAT2, among others) is certainly a contributory factor to gene expression changes associated with β cell failure^13-15^, recent studies have linked long non-coding RNAs (lncRNAs) to the regulation of innate immune responses in different cell types^16-18^. LncRNAs are non-coding RNAs with a length of 200 nucleotides or more with a structure similar to the one of protein-coding genes^19^. Several studies have implicated lncRNAs in biological and cellular processes related to inflammation, including activation of the innate antiviral immune response through the activation of Pattern Recognition Receptor (PRR)-related signal transduction^20^, regulation of innate immune-associated chemokines^21^ and other inflammatory genes^22^. The function of non-coding RNAs is only beginning to emerge but there is already strong evidence of their involvement in disease, including inflammatory and autoimmune disorders^5,23-25^.

Genome-wide association studies (GWAS) have identified a large number of single nucleotide polymorphisms (SNPs) predisposing to immune diseases^26^. Only a small fraction of these SNPs is located within protein-coding genes and the majority map to non-coding regions of the human genome^27^. Around 10% of the SNPs associated to immune disorders lie in lncRNAs^28^, suggesting that risk variants in these non-coding molecules could alter their function and dysregulate gene expression networks potentially important for pancreatic β cell function and T1D development.

A combination of expression data and DNA sequence variation in T1D previously led to the description of an antiviral gene expression network linked to T1D susceptibility named IDIN (IRF7-driven inflammatory network)^29^. In this study, Heinig et al showed that the genotype of an intergenic T1D risk SNP (rs9585056) correlated with the expression of the IDIN network in immune cells. In the current study, we show that rs9585056 is not intergenic but is located in NONHSAT233405.1 or *ARGI* (Antiviral Response Gene Inducer), a lncRNA gene that is upregulated by Coxsackievirus infections in pancreatic β cells.

We functionally characterized *ARGI* and demonstrate that it regulates the expression of antiviral and pro-inflammatory genes in pancreatic β cells. *ARGI* participates in pancreatic β cell inflammation via transcriptional regulation of interferon stimulated genes (ISGs). Since type I IFN signaling plays a crucial role in T1D-related β cell dysfunction^2^, *ARGI* may be functionally implicated in the pathogenesis of the disease. These results serve as a proof of concept of the potential implication of T1D-associated lncRNAs in the dysfunction of pancreatic β cells in T1D, and open a new avenue for the development of therapeutic approaches based on lncRNA expression modification.

## Results

### ARGI is a nuclear lncRNA upregulated in EndoC-βH1 cells upon a viral infection

To identify potential lncRNAs associated with T1D, the genomic location of all T1D-associated SNPs (GWAS catalog; https://www.ebi.ac.uk/gwas) were intersected with the genomic location of all lncRNAs annotated in NONCODE version 6 (http://www.noncode.org/index.php). Through this approach, we identified 69 lncRNAs harboring at least one SNP associated with the disease (Fig EV1). Of those 69, 19 lncRNAs have a T1D-associated SNP in their exonic region. Because SNPs in lncRNAs may affect their function through the disruption of their secondary structure^5,25^, we assessed whether these 19 SNPs alter lncRNA secondary structure. *In silico* predictions using RNAsnp software (Center for non-coding RNA in Technology and Health) revealed that the secondary structure of 10 lncRNAs was altered by at least one of the T1D-associated SNPs.

We next examined whether these 10 T1D-associated lncRNAs were expressed in pancreatic β cells and modulated by viral infections. To this end, we analyzed their expression in the human β cell line EndoC-βH1 in basal condition and after transfection with polyinosinic-polycytidylic acid (PIC), a molecule that simulates viral dsRNA produced during viral infections. Nine out of the analyzed 10 lncRNAs were expressed in pancreatic β cells, and as shown in Fig 1A, all except *LncRNA_7* were upregulated 8 and/or 24 h after PIC transfection.

**Figure 1.**
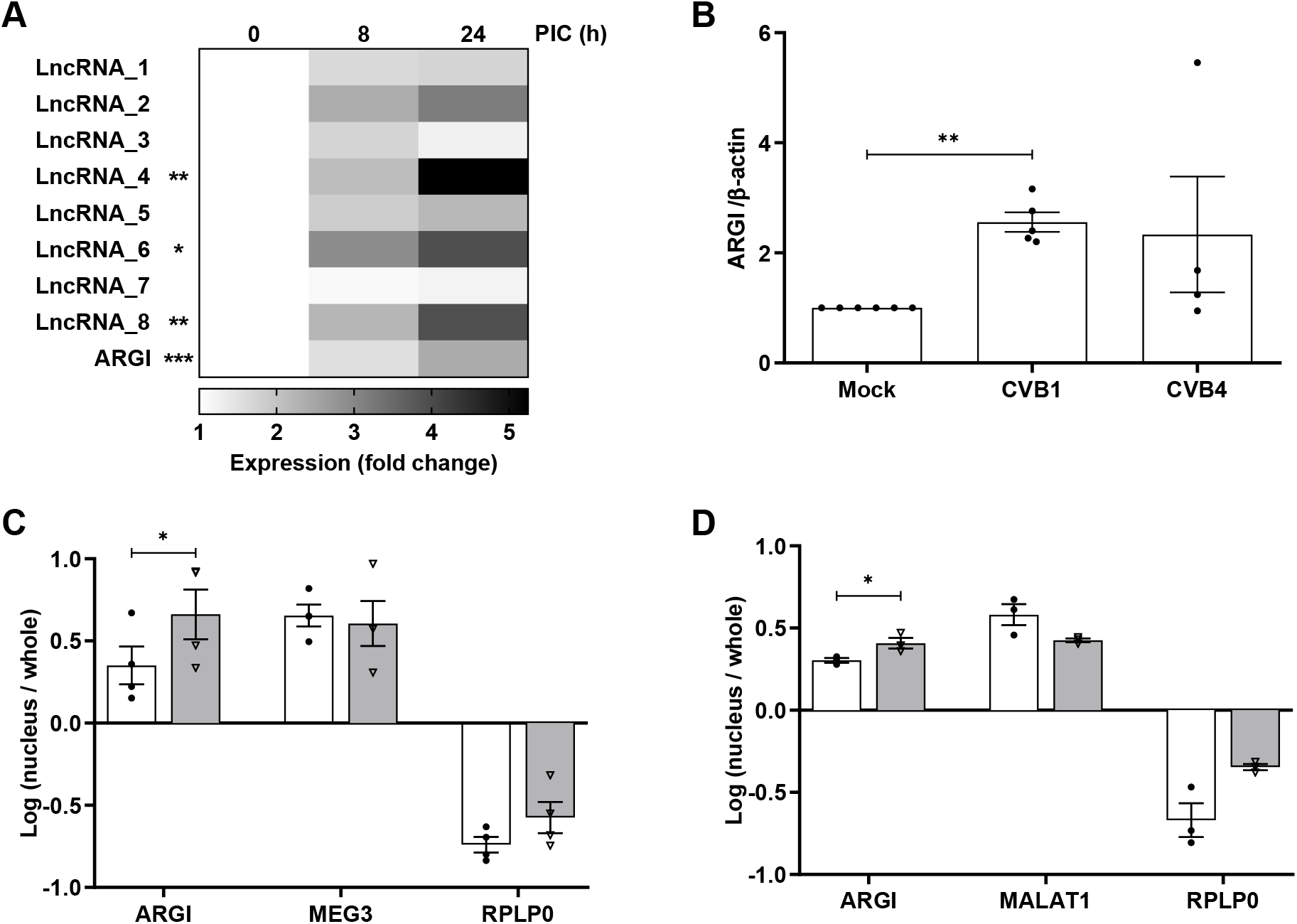
*ARGI* is a nuclear lncRNA upregulated in EndoC-βH1 cells upon viral infection. (A) EndoC-βH1 cells were exposed to intracellular PIC (1 μg/mL) for 8 or 24h and the expression of ten T1D-associated lncRNAs was determined by qPCR. One of ten lncRNAs was not detected and the expression of the other 9 lncRNAs in EndoC-βH1 cells is presented as a heatmap (fold change vs. PIC 0h). Results are means of 4 independent experiments; *p< 0.05, **p<0.01, ***p<0.001; Student’s t test. (B) Induced pluripotent stem cell (iPSC)-derived pancreatic β-like cells were left uninfected (Mock) or infected with CVB1 or CVB4 for 24h. *ARGI* expression was assessed by qPCR and normalized to the reference gene β-actin. Data are means±SEM of 5 to 6 independent iPSC differentiations; **p<0.01; Student’s t test. (C) EndoC-βH1 cells were left untransfected (white bars) or exposed to PIC (1 μg/mL) for 24h (grey bars). Relative *ARGI* expression was determined in nucleus and whole cell extracts, using *MEG3* and *RPLP0* as controls for the respective fractions. Amounts of specific nuclear RNA were measured by qPCR and compared to the total amount of RNA in the whole cell extract. Data are expressed as a logarithm (nucleus/whole) and are means±SEM of 4 independent experiments; *p< 0.05; Student’s t test. (D) EndoC-βH1 cells were left uninfected (white bars) or infected with CVB1 (MOI 0.05, grey bars). Relative *ARGI* expression was determined in nucleus and whole cell extracts, using *MALAT* and *RPLP0* as controls for nuclear and whole cell fractions, respectively. Amounts of specific nuclear RNA were measured by qPCR and compared to the total amount of RNA in the whole cell. Data are expressed as a logarithm (nucleus/whole) and are means±SEM of 3 independent experiments; *p< 0.05; Student’s t test.

One of the PIC-upregulated T1D-associated lncRNAs, *ARGI* (Antiviral Response Gene Inducer), harbors a SNP (rs9585056) in its third exon that was previously described as an intergenic SNP that has an eQTL regulating expression of the IDIN antiviral gene network in monocytes and macrophages^29^. Our mapping of T1D-associated SNPs against lncRNAs showed, however, that this SNP is not intergenic but falls into the hitherto uncharacterized lncRNA *ARGI*. Moreover, our *in silico* prediction suggested that the SNP disrupts *ARGI*’s secondary structure by affecting one of its loops (Fig EV2). Considering that enteroviral infections such as coxsackievirus B (CVB) may contribute to T1D pathogenesis by infecting pancreatic β cells^12,30-32^, we set out to characterize the function of *ARGI* at the pancreatic β cell level.

We first examined whether CVB infections modify *ARGI* expression in pancreatic β cells. To this end, we differentiated human induced pluripotent stem cells (iPSCs) into pancreatic β cells using a 7-stage method that mimics embryonic β cell development. At the end of the differentiation, the iPSC-derived aggregates contained 47.9±3.48% β cells, 2.7±1.22% α cells and 0.7±0.08% δ cells. We infected these human iPSC-derived β cells with two diabetogenic CVB strains, namely CVB1 and CVB4. CVB1 increased *ARGI* expression, while the induction with CVB4 was more variable (Fig 1B).

Considering that cellular location often determines lncRNA function^19,33-35^, we next examined the subcellular localization of *ARGI* in the human β cell line EndoC-βH1. In basal condition, *ARGI* was preferentially expressed in the nucleus of EndoC-βH1 cells (Fig 1C), and its nuclear expression was upregulated after 24-h PIC exposure. In keeping with the results with this viral dsRNA mimic, CVB1 infection induced nuclear *ARGI* expression in EndoC-βH1 cells (Fig 1D).

### ARGI upregulation in pancreatic β cells leads to hyperactivation of an inflammatory and antiviral gene signature

In order to dissect the biological impact of *ARGI* upregulation in response to viral dsRNA (PIC), we performed RNA-sequencing of *ARGI*-overexpressing pancreatic β cells. *ARGI* was upregulated in the EndoC-βH1 cell line using an overexpression vector (Fig 2A). Interestingly, Gene Ontology (GO) enrichment analysis revealed that the upregulated genes were enriched in pathways related to antiviral responses and type I IFN signaling (cellular response to type I IFNs, type I IFN signaling pathway, defense response to virus, among others) (Fig 2B).

**Figure 2.**
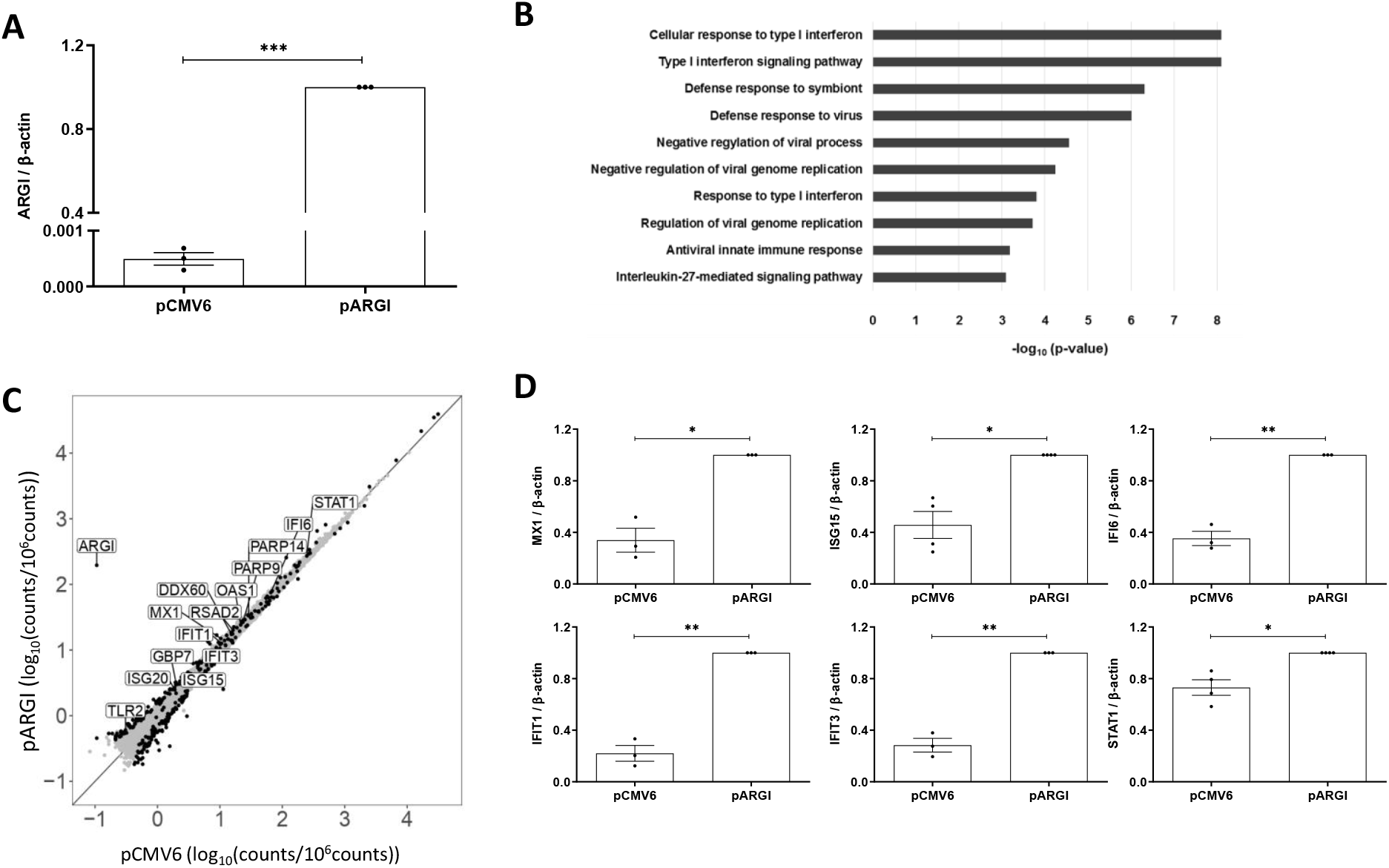
*ARGI* upregulation in pancreatic β cells induces an inflammatory and antiviral gene signature. (A) EndoC-βH1 cells were transfected with an empty overexpression plasmid (pCMV6) or with a plasmid overexpressing *ARGI* (pARGI). *ARGI* expression was determined by qPCR and normalized to the reference gene β-actin. Results are means±SEM of 3 independent experiments; ***p<0.001; Student’s t test. (B) Gene Ontology analysis shows enrichment of inflammatory and antiviral pathways in *ARGI*-overexpressing pancreatic β cells. (C) Scatterplot showing differentially expressed genes (black dots) in *ARGI*-overexpressing EndoC-βH1 cells (pARGI) compared to pCMV6-transfected control cells. Expression values are presented as Log_10_ (counts/10^6^ counts). *ARGI* and genes from the IDIN network are indicated. (D) EndoC-βH1 cells were transfected with a control empty plasmid (pCMV6) or with a *ARGI* plasmid (pARGI). Expression of *MX1, ISG15, IFI6, IFIT1, IFIT3* and *STAT1* genes was determined by qPCR and normalized to the reference gene β-actin. Results are means±SEM of 3 independent experiments; **p < 0.01 and *p < 0.05; Student’s t test.

Among the upregulated genes, members of the IDIN gene expression network were significantly overrepresented (Fig 2C). Among the 17,249 transcripts detected by RNA-seq, 430 (2.5%) were genes of the IDIN network, and 4.3% of the significantly upregulated genes were IDIN members (p-value: 0.0188). The top upregulated IDIN genes included the ISGs *MX1, ISG15, IFI6, IFIT1, IFIT3* and *STAT1*. Expression of these top six upregulated genes in *ARGI*-overexpressing beta cells was confirmed in an independent EndoC-βH1 cell sample set by qPCR (Fig 2D).

### ARGI participates in the regulation of virus-induced IFNβ and ISG expression in pancreatic β cells

Because most of the *ARGI*-upregulated IDIN genes were ISGs^36^, we asked whether *IFNβ* expression was induced in *ARGI*-overexpressing β cells. *ARGI* overexpression increased *IFNβ* expression by 60% in pancreatic β cells (Fig 3A), suggesting that *ARGI* might, at least in part, regulate ISGs through *IFNβ* induction.

**Figure 3.**
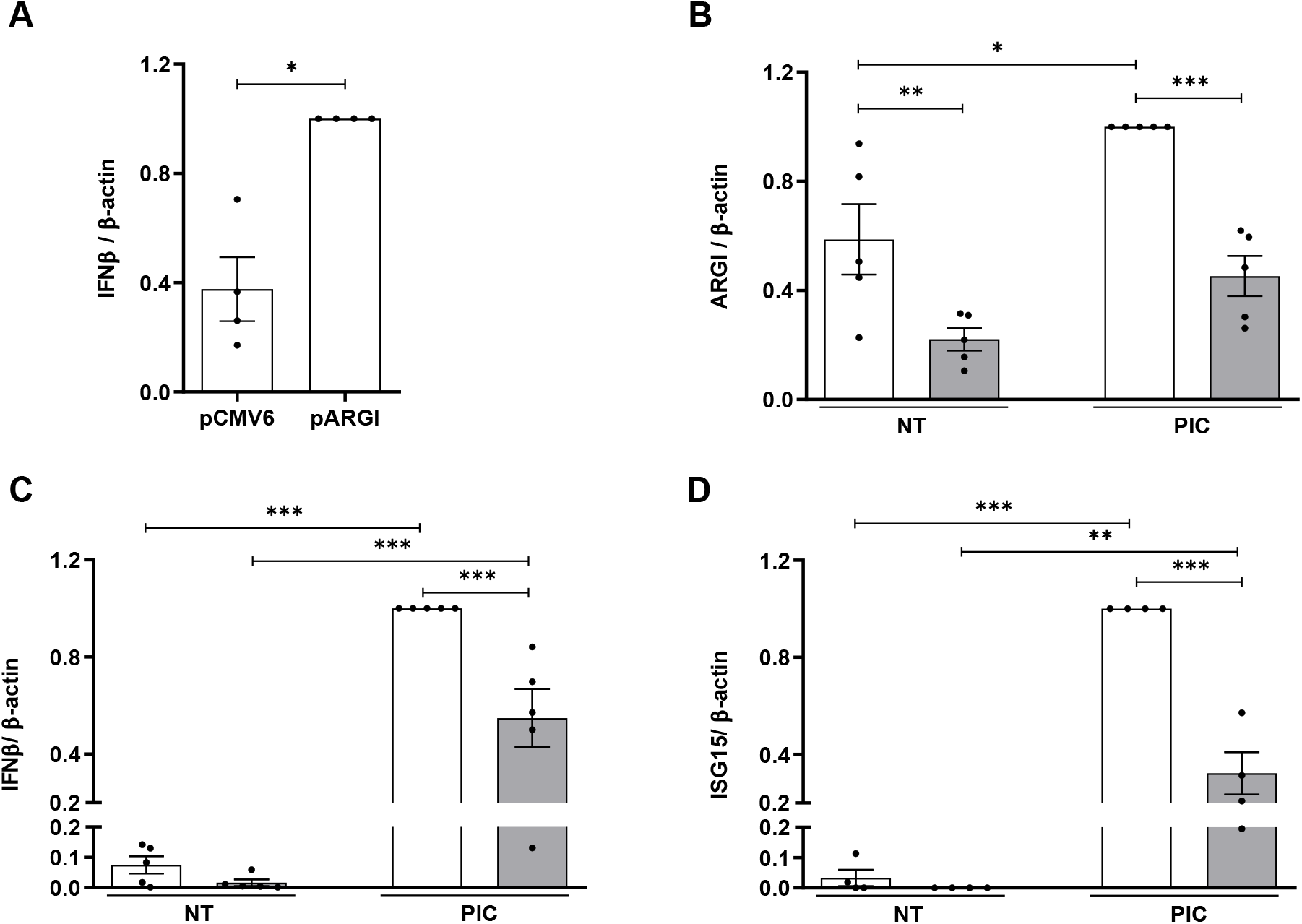
*ARGI* participates in the regulation of virus-induced *IFNβ* and ISG expression in pancreatic *β* cells. (A) EndoC-βH1 cells were transfected with pCMV6 or pARGI. *IFN*β expression was determined by qPCR and normalized to the reference β-actin. Results are means±SEM of 4 independent experiments. *p < 0.05; Student’s t test (B) Transcriptional activation of *ARGI* was inhibited using CRISPRi. EndoC-βH1 cells were transfected with an empty CRISPRi vector (white bars) or with a CRISPRi vector harboring a sgRNA targeting a conserved NFκB binding site in *ARGI*’s regulatory region (grey bars). After 36h of transfection, cells were left non-transfected (NT) or transfected with PIC (1μg/ml) for 24h. Expression of *ARGI* (B), *IFNβ* (C) and *ISG15* (D) was determined by qPCR and normalized to the reference gene β-actin. Results are means±SEM of 4-5 independent experiments; ***p < 0.001, **p < 0.01 and *p < 0.05 as indicated; Student’s t test or ANOVA followed by Student’s t test.

To test whether *ARGI* is implicated in IFNβ transcription, we deleted *ARGI* by CRISPR-Cas9 in pancreatic β cells (Fig EV3A). Due to the low replication rate of EndoC-βH1 cells, we could not select a complete knockout clone. In basal condition, *ARGI* expression was similar in CRISPR-Cas9-transfected and control cells; in cells exposed to intracellular PIC, however, *ARGI* expression was 30-50% lower in CRISPR-Cas9-transfected cells (Fig EV3B). This led to a 60-70% decrease in PIC-induced *IFNβ* and *ISG15* expression in pancreatic β cells (Fig S3C-D).

To confirm these results, we inhibited endogenous *ARGI* expression by CRISPRi technique using a sgRNA targeting a conserved NFκB binding site in a potential regulatory region close to the *ARGI* transcription starting site (Fig EV4A). We first confirmed that *ARGI* was partially regulated by NFκB using the specific chemical inhibitor BAY 11-7082. As shown in Figure EV4B, PIC-induced *ARGI* upregulation was counteracted by NFκB inhibition. Using the CRISPRi-sgRNA, *ARGI* expression was downregulated by around 50% both in basal and PIC-transfected conditions (Fig 3B). This decreased in PIC-induced *IFNβ* expression by 40% (Fig 3C). In keeping with the CRISPR-Cas9 experiment, PIC-induced *ISG15* expression was also reduced upon *ARGI* inhibition (Fig 3D). These results show that *ARGI* regulates expression of *IFNβ* as well as other ISGs, such as *ISG15*.

### *ARGI* associates with the transcription factor CTCF to bind to the regulatory regions of IFNβ and *ISG15* genes in PIC-transfected cells

In order to determine the molecular mechanisms by which *ARGI* regulates expression of *IFNβ* and ISGs, we next performed an RNA antisense purification experiment to purify *ARGI*-bound chromatin in non-treated and PIC-treated pancreatic β cells, and analyzed whether the regulatory regions of *IFNβ* and *ISG15* were present by qPCR. *ARGI* was purified using biotinylated antisense complementary oligonucleotides. Antisense oligonucleotides complementary to an unrelated lncRNA of similar size were used as negative control (Fig 4A and Fig EV5A). *ARGI* purification was more efficient in PIC-transfected cells than in non-transfected cells, most probably due to *ARGI* induction by PIC (Fig 4A). The binding of *ARGI* to *IFNβ* and *ISG15* regulatory regions was determined by qPCR using primer pairs targeting promoter and enhancer regions of *IFNβ* and *ISG15* genes (Fig EV5B-C). As shown in Fig 4B-D, in basal condition, *ARGI* bound *IFNβ* promoter but it did not bind the *ISG15* promoter or *ISG15* enhancer. In PIC-treated β cells, *IFNβ* promoter, *ISG15* promoter and *ISG15* enhancer levels were increased both, compared with non-PIC-transfected β cells and relative to PIC-transfected cells transfected with an unrelated control lncRNA.

**Figure 4.**
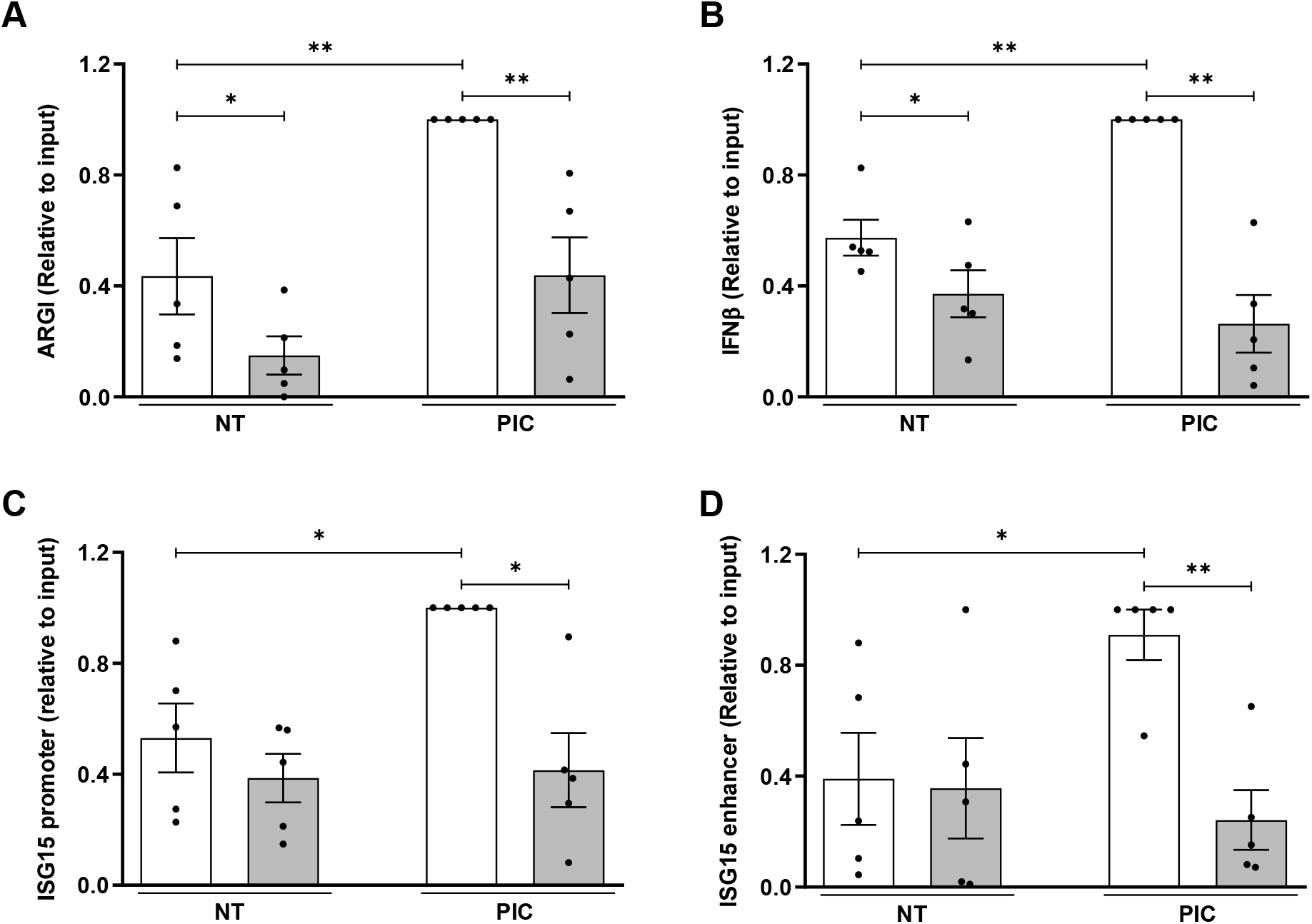
*ARGI* binds to *IFNβ* and *ISG15* regulatory regions upon viral insult. RNA antisense purification of *ARGI* was performed in non-transfected (NT) or PIC-transfected EndoC-βH1 cells (PIC). (A) *ARGI* was purified using biotinylated antisense complementary oligonucleotides (white bars); antisense oligonucleotides complementary to an unrelated similar length lncRNA were used as negative control (grey bars). (B-D) *ARGI*-bound *IFNβ* promoter (B), *ISG15* promoter (C) and *ISG15* enhancer (D) amounts were determined by qPCR. Results are expressed as relative to input and are means±SEM of 5 independent experiments. **p<0.01 and *p < 0.05 as indicated; ANOVA followed by Student’s t test.

The regulatory regions (promoters and enhancers) of *IFNβ* and *ISG15* contain binding sites for several key pro-inflammatory transcription factors (IRF7, STAT1 and STAT2, among others). In addition, there are binding sites for CCCTC-binding factor (CTCF), a conserved zinc finger protein that can act as a transcriptional activator, repressor or insulator protein, blocking the communication between enhancers and promoters^37,38^. *In silico* prediction of RNA-protein interactions using the CatRAPID tool^39^ revealed that *ARGI* potentially interacts with CTCF (interaction score: 0.41), but not with STAT1, STAT2 or IRF7 proteins. To experimentally assess whether *ARGI* interacts with CTCF, we performed RNA immunoprecipitation using a CTCF targeting antibody in basal and PIC-transfected pancreatic β cells (Fig 5A). Under basal conditions, *ARGI* and CTCF did not interact, but upon exposure to intracellular PIC, *ARGI* bound to CTCF in pancreatic β cells (Fig 5B).

**Figure 5.**
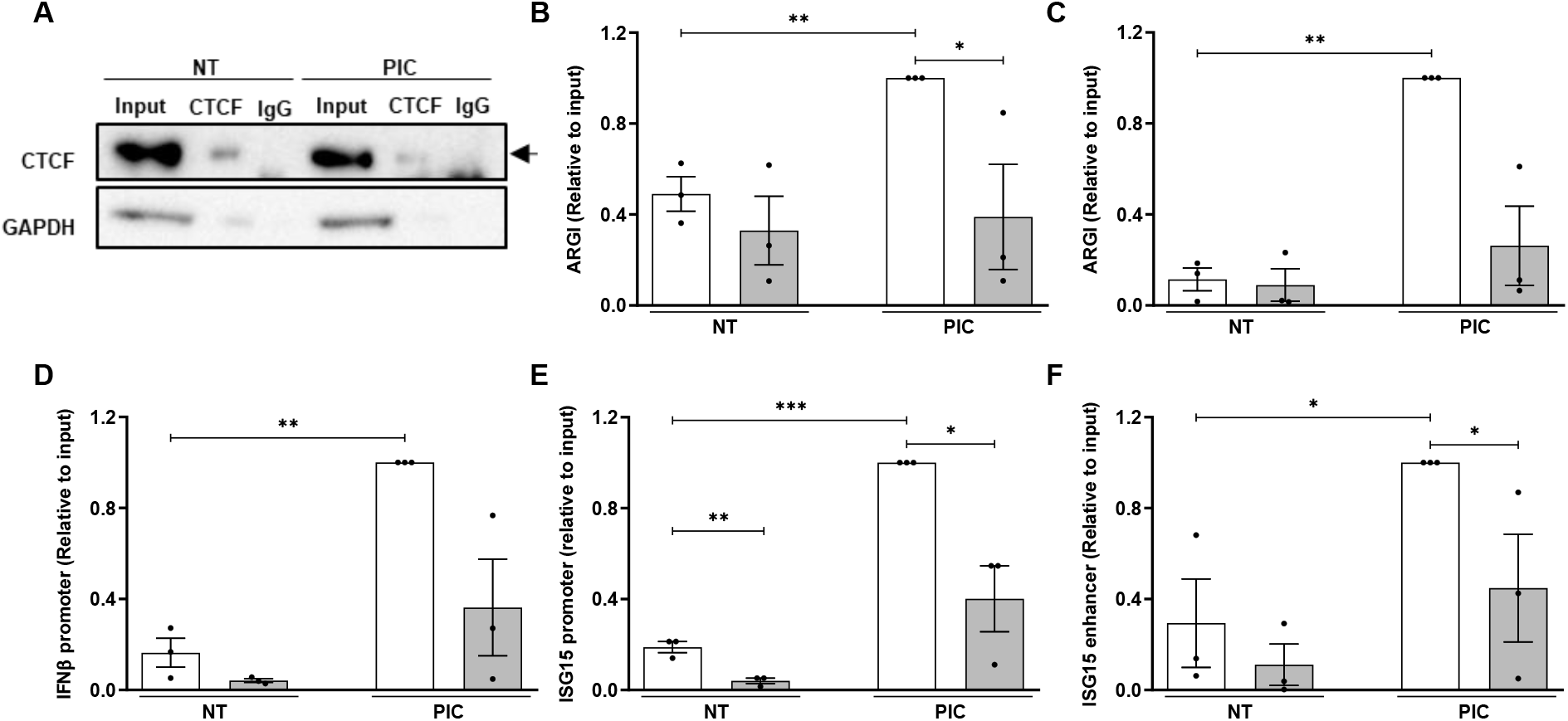
*ARGI* associates with the transcription factor CTCF to bind the regulatory regions of *IFNβ* and *ISG15* genes in viral dsRNA-transfected cells. (A) EndoC-βH1 cells were left non-transfected (NT) or transfected with PIC for 24h and RNA immunoprecipitation was performed using a CTCF antibody or an IgG antibody used as a negative control. The image is representative of 3 independent experiments. (B) *ARGI* expression was determined in CTCF-bound RNA (white bars) or IgG-bound RNA (grey bars) by qPCR. Results are means±SEM of 3 independent experiments and the amounts of *ARGI* are expressed as relative to input. **p<0.01, and *p<0.05 as indicated; ANOVA followed by Student’s t test. (C-F) Chromatin RNA immunoprecipitation was performed in non-transfected (NT) and PIC-transfected HEK293 cells. CTCF-bound chromatin and RNA were immunoprecipitated using an antibody for CTCF (white) or IgG (grey), used as negative control. CTCF-bound *ARGI* expression (C), *IFNβ* promoter (D), *ISG15* promoter (E) and *ISG15* enhancer amounts (F) were determined by qPCR. Results are means±SEM of 3 independent experiments and are presented as relative to the input. ***p<0.001, **p<0.01, and *p<0.05 as indicated; ANOVA followed by Student’s t test.

To check whether *ARGI* and CTCF in turn interact with *IFNβ* and *ISG15* regulatory regions, we next performed a chromatin-RNA immunoprecipitation to immunoprecipitate RNA and chromatin fragments simultaneously bound to CTCF. To this end, CTCF-bound chromatin and RNA were captured using an anti-CTFC antibody, and *ARGI, IFNβ* promoter and *ISG15* promoter and enhancer were amplified by qPCR. As shown in Fig 5C-F, CTCF was bound to *ARGI* and simultaneously interacted with *IFNβ* promoter, *ISG15* promoter and enhancer, especially upon PIC transfection.

### Allele-specific binding of *ARGI* to CTCF affects IDIN gene expression levels

LncRNA secondary structure is crucial for its function since it may affect its stability and binding to DNA, proteins or other RNAs^40-43^. *ARGI* harbors a T1D-associated SNP that was predicted to affect its secondary structure (Fig EV2). To test whether the T1D-associated SNP affects *ARGI* function, we first determined the interaction of *ARGI* and CTCF in the presence of the T1D protective or risk allele. To this end, we performed RNA immunoprecipitation in cells overexpressing CTCF and *ARGI* harboring the T1D protective (ARGI-P) or risk allele (ARGI-R) (Fig 6A). As shown in Fig 6B, *ARGI* containing either allele interacted with CTCF, but the interaction was stronger when *ARGI* harbored the T1D risk allele (rs9585056-G) than when it had the protective allele (rs9585056-A).

**Figure 6.**
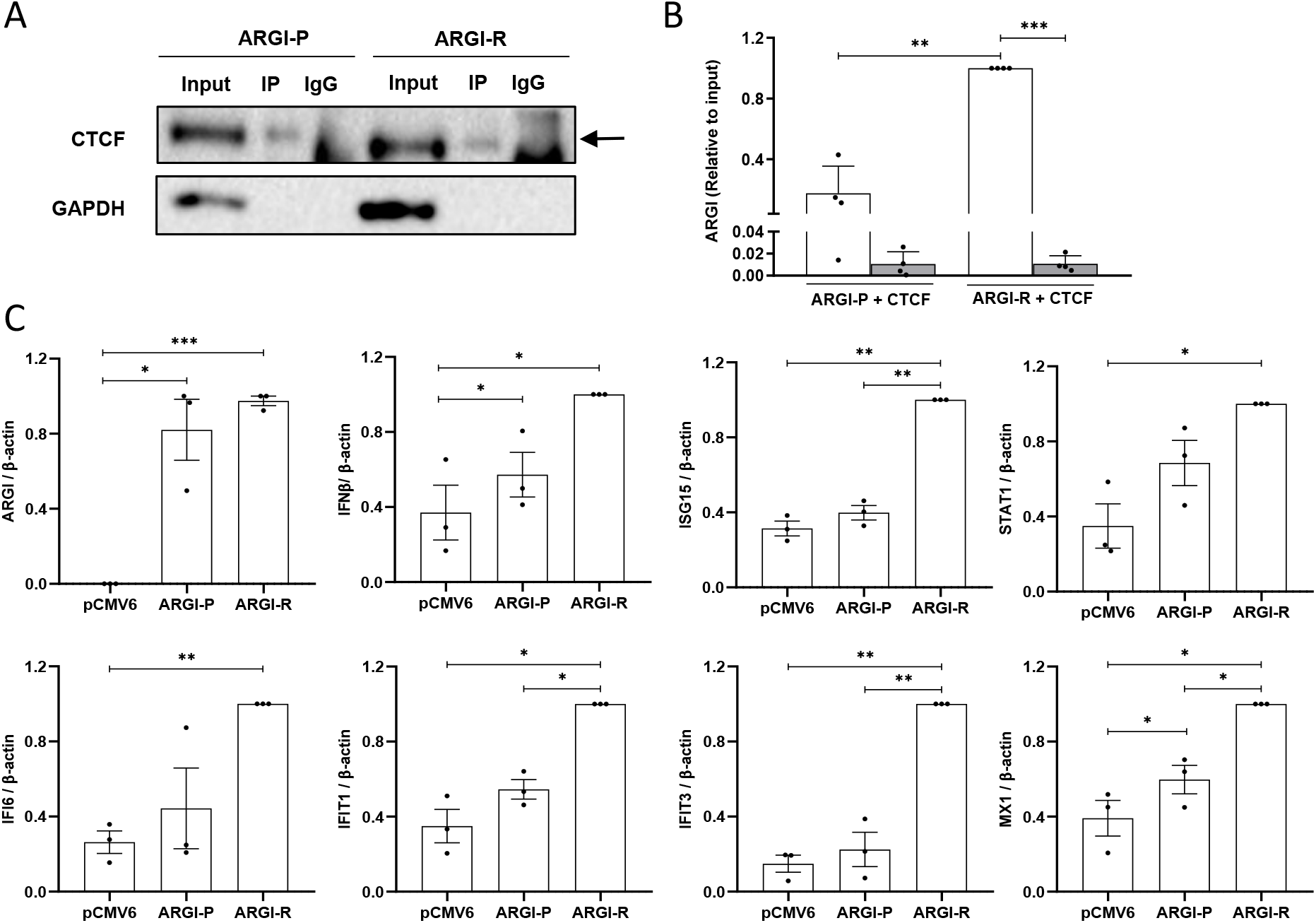
Allele-specific binding of *ARGI* to CTCF affects IDIN gene expression. (A) EndoC-βH1 cells were transfected with a plasmid overexpressing *ARGI* harboring the T1D protective (ARGI-P) or risk allele (ARGI-R) and co-transfected with a vector overexpressing CTCF. RNA immunoprecipitation was performed using an antibody for CTCF; IgG was used as negative control. (B) CTCF-(white bars) and IgG-bound (grey bars) *ARGI* amounts were determined by qPCR. Results are means±SEM of 4 independent experiments and the amounts of *ARGI* are presented as relative to the input. ***p<0.001 and **p<0.01 as indicated; ANOVA followed by Student’s t test. (C) EndoC-βH1 cells were transfected with control plasmid (pCMV6) or plasmids overexpressing *ARGI* harboring the protective (ARGI-P) or risk allele (ARGI-R). Expression of *ARGI, MX1, ISG15, STAT1, IFI6, IFIT1, IFIT3* and *IFNβ* was determined by qPCR and normalized to the reference gene β-actin. Results are means±SEM of 3 independent experiments and are presented as relative to the input. ***p<0.001, **p<0.01, and *p<0.05 as indicated; ANOVA followed by Student’s t test.

We next analyzed the expression of *IFNβ, ISG15* and other IDIN genes in EndoC-βH1 cells overexpressing ARGI-P or ARGI-R (Fig 6C). Both vectors induced similar *ARGI* expression, suggesting that the T1D-associated SNP did not affect *ARGI* stability. Interestingly, overexpression of *ARGI* harboring the T1D risk allele induced a higher expression of *IFNβ, ISG15, MX1, STAT1, IFI6, IFIT1* and *IFIT3* genes than overexpression of *ARGI* harboring the T1D protective allele (Fig 6C).

## Discussion

T1D is a complex autoimmune disease in which genetic and environmental factors interact to trigger an autoimmune assault against pancreatic β cells^2^. Several *loci* throughout the human genome have been associated with genetic risk for T1D, and several candidate genes have been proposed as being causal. Many of the T1D candidate genes characterized so far have been implicated in the regulation of antiviral and pro-inflammatory responses at the pancreatic β cell level^44^. Some participate in the regulation of the type I IFN signaling, such as *PTPN2* and *TYK2*, by regulating the type I IFN-induced JAK/STAT signaling pathway^3,4,45^, while others (e.g. *MDA5*) encode viral dsRNA cytoplasmic receptors^6^. The interaction between genetic variants in T1D candidate genes and viral infections have been studied these past years, demonstrating that T1D risk alleles hyperactivate antiviral and pro-inflammatory responses in pancreatic β cells, that eventually lead to β cell destruction and T1D development^45-48^.

Over the past years, advances in the annotation of the human genome have revealed that many disease-associated SNPs are located within lncRNAs, affecting their function by disrupting their secondary structure^24^. For instance, the T1D-associated lncRNA *Lnc13* regulates virus-induced STAT1 signaling in pancreatic β cells in an allele-specific manner^5^. The implication of lncRNAs in the regulation of pro-inflammatory pathways has also been documented in other contexts. For example, the lncRNA *LUCAT1* is a negative feedback regulator of IFN responses in humans by interacting with STAT1 in the nucleus^49^, and lncRNA *Mirt2* inhibits activation of NFκB and MAPK pathways, limiting the reduction of LPS-induced proinflammatory cytokines, through attenuation of Lys63 (K63)-linked ubiquitination of TRAF6 in macrophages^50^.

Herein, we functionally characterized the lncRNA *ARGI* (Antiviral Response Gene Inducer) in the regulation of virus-induced pancreatic β cell inflammation. The SNP associated with T1D (rs9585056) was initially catalogued as an intergenic SNP that correlated with the expression of the antiviral gene network IDIN^29^. IDIN is enriched with ISGs, such as *ISG15, MX1* or *IF6*, among others. Our analysis showed that the T1D-associated rs9585056 is actually located in an exon of *ARGI*, and *in silico* predictions revealed that the SNP disrupts its secondary structure.

RNA-sequencing experiments of *ARGI*-overexpressing pancreatic β cells revealed a role for *ARGI* in antiviral and pro-inflammatory gene expression regulation, with upregulation of type I IFN signaling and antiviral pathways. IDIN genes were significantly overrepresented among the genes induced by *ARGI*-overexpression, suggesting that *ARGI* may regulate IDIN in pancreatic β cells.

Importantly, our data showed that allele-specific upregulation of *ARGI* in pancreatic β cells led to increased expression of IDIN genes in an allele-specific manner. *ARGI* harboring the T1D risk allele induced a higher expression of IDIN genes than the lncRNA harboring the T1D protective allele. Since the secondary/tertiary structure of lncRNAs seems to be crucial for their function, genetic variants and mutations potentially contribute to disease pathogenesis by altering disease-associated pathways^5,24,51^. We found that *ARGI* regulates the expression of some ISGs in an allele-specific manner through its interaction with the transcription factor CTCF. Viral dsRNA triggered translocation of *ARGI* to the nucleus of pancreatic β cells. Once in the nucleus, *ARGI* interacts with CTCF to bind to regulatory regions of some ISGs - e.g. *IFN*β and *ISG15* - to transcriptionally activate them. Also under basal conditions, the interaction between CTCF and *ARGI* is stronger for the T1D risk allele than for the protective allele, suggesting that stronger binding between *ARGI* and CTCF will exacerbate ISG expression in pancreatic β cells. Other studies have shown that interaction between disease-associated lncRNAs and transcription factors, both at the mRNA and protein level, may affect gene expression in an allele-specific manner. Indeed, *Lnc13* in pancreatic β cells interacts in allele-specific manner with PCBP2 to stabilize *STAT1* mRNA, leading to increased STAT1 activation and pro-inflammatory chemokine expression when the T1D risk allele is present in *Lnc13*^5^. Along the same lines, a lncRNA harboring a SNP associated with cancer (*CCAT2*) regulates cancer cell metabolism through its interaction with the Cleavage factor I (CFIm) in an allele-specific manner^52^.

The transcription factor CTCF is a ubiquitously expressed multifunctional transcriptional regulator. It can inhibit or activate gene transcription depending on the cell type, location of the binding site, stimulus and presence of interacting partners (transcriptional activators, repressors, cohesins, and RNA pol II^53^. Using RNA antisense purification and RNA immunoprecipitation, we observed that in the presence of viral dsRNA, *ARGI* and CTCF interact with the regulatory regions of *IFNβ* and *ISG15*, suggesting that viral infections in β cells promote ISG expression through the binding of *ARGI* and CTCF to regulatory regions of these antiviral genes. Recent studies have shown that CTCF can bind non-coding RNAs to modulate gene expression, including inflammation-related genes^54,55^. Using RedCHIP technique, a recent study identified several of cis- and trans-acting non-coding RNAs enriched at CTCF-binding sites in human cells, which may be involved in in CTCF-dependent chromatin looping and gene activation/repression^54^. Some studies have characterized the impact of CTCF and non-coding RNA interaction in the regulation of inflammatory genes. For example, a non-coding RNA derived from normally silenced pericentromeric repetitive sequences was shown to impair CTCF binding to chromatin, resulting in the alteration of chromatin accessibility and the activation of SASP-like inflammatory genes^55^.

In conclusion, our data demonstrate that *ARGI* is implicated in the regulation of virus-induced inflammation in pancreatic β cells through a molecular mechanism that implies allele-specific binding to the transcription factor CTCF and regulation of ISG expression, a gene expression signature related to the T1D pathogenesis (Fig 7). Our results are in concordance with the expression of type I IFNs in pancreatic islets of T1D donors^56-58^. Moreover, more recent studies, such as the Diabetes Virus Detection (DiViD) study, have observed upregulation of several ISGs, including *STAT1, IFI6* or *MX1* in islets from individuals with T1D^48,59^.

**Figure 7.**
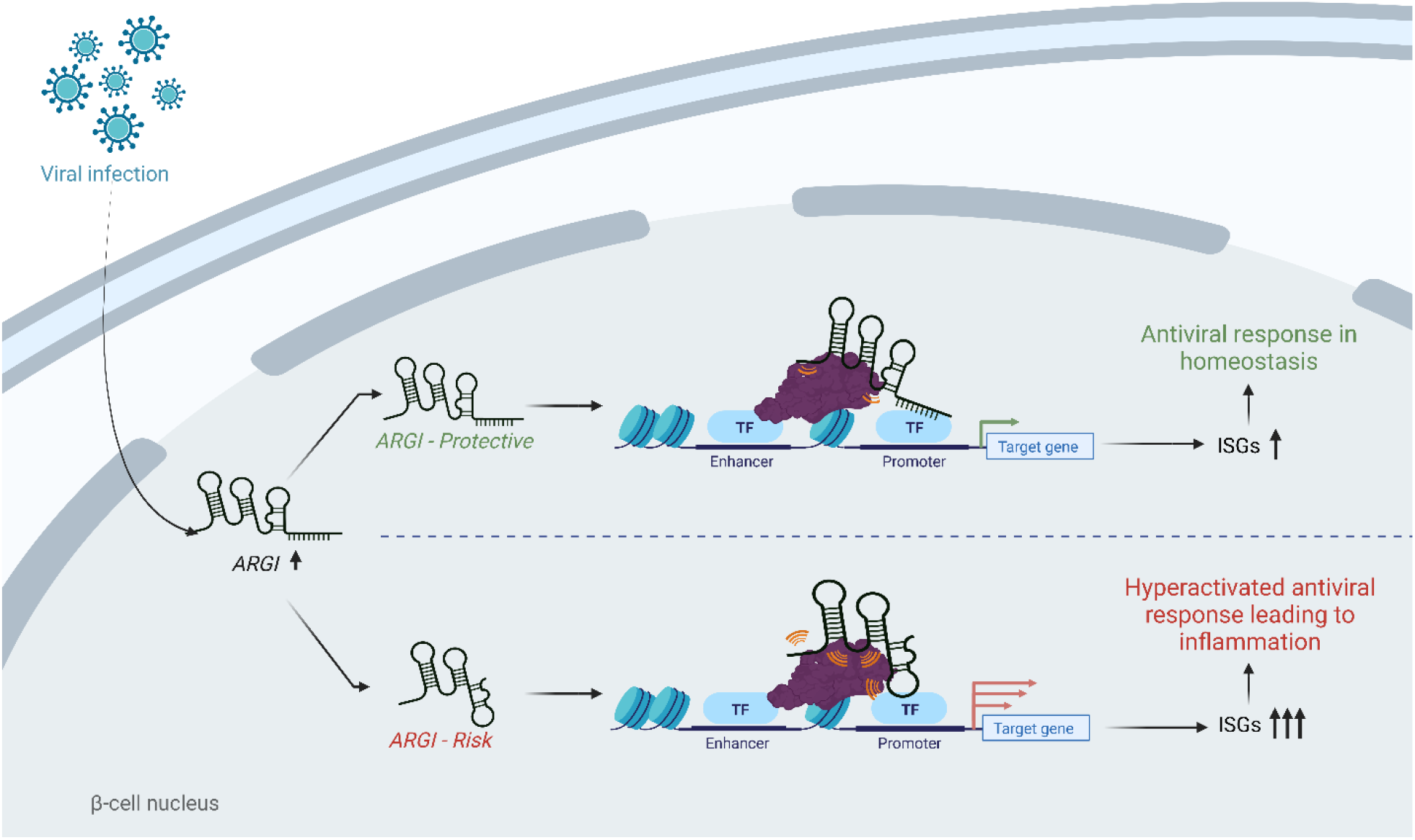
A model of the role of *ARGI* in the allele-specific modulation of virus-induced ISG transcription in pancreatic beta cells. Viral infections trigger *ARGI* upregulation in the nuclei of pancreatic β cells. *ARGI* interacts with the transcription factor CTCF to bind to the regulatory regions of interferon stimulated genes (ISGs), promoting their transcription. When *ARGI* harbors the T1D protective allele, a homeostatic antiviral response is activated; however when the T1D risk allele is present, this process is exacerbated, leading to an hyperactivation of the antiviral response in pancreatic β cells. In the context of T1D pathogenesis, the generation of an excessive pro-inflammatory environment in pancreatic islets, leads to increased insulitis, and eventually, to pancreatic β cell destruction and T1D development.

The implication of viral infections and the resulting activation of type I IFN signaling in β cells seems to be crucial in the initial stages of T1D, however the specific pathogenic mechanisms that trigger the infection-induced autoimmune assault against β cells are not fully elucidated. Accumulating evidence suggest that the factor linking viral infections with the autoimmune triggering rely on the presence of T1D-associated genetic variants in genes expressed in pancreatic β cells^3-6^. Hence, the functional characterization of T1D susceptibility genes in virus-induced pancreatic β cell dysfunction will help to clarify many of the “unknowns” that still linger regarding the pathogenesis of the disease. In this sense, our findings provide novel information on the molecular mechanisms by which disease-associated SNPs in lncRNAs influence the activation of diabetogenic gene expression pathways in pancreatic β cells, and on how interactions between T1D-associated genes and infections may trigger inflammation in the initial stages of the disease.

## Materials and Methods

### Reagents and tools table

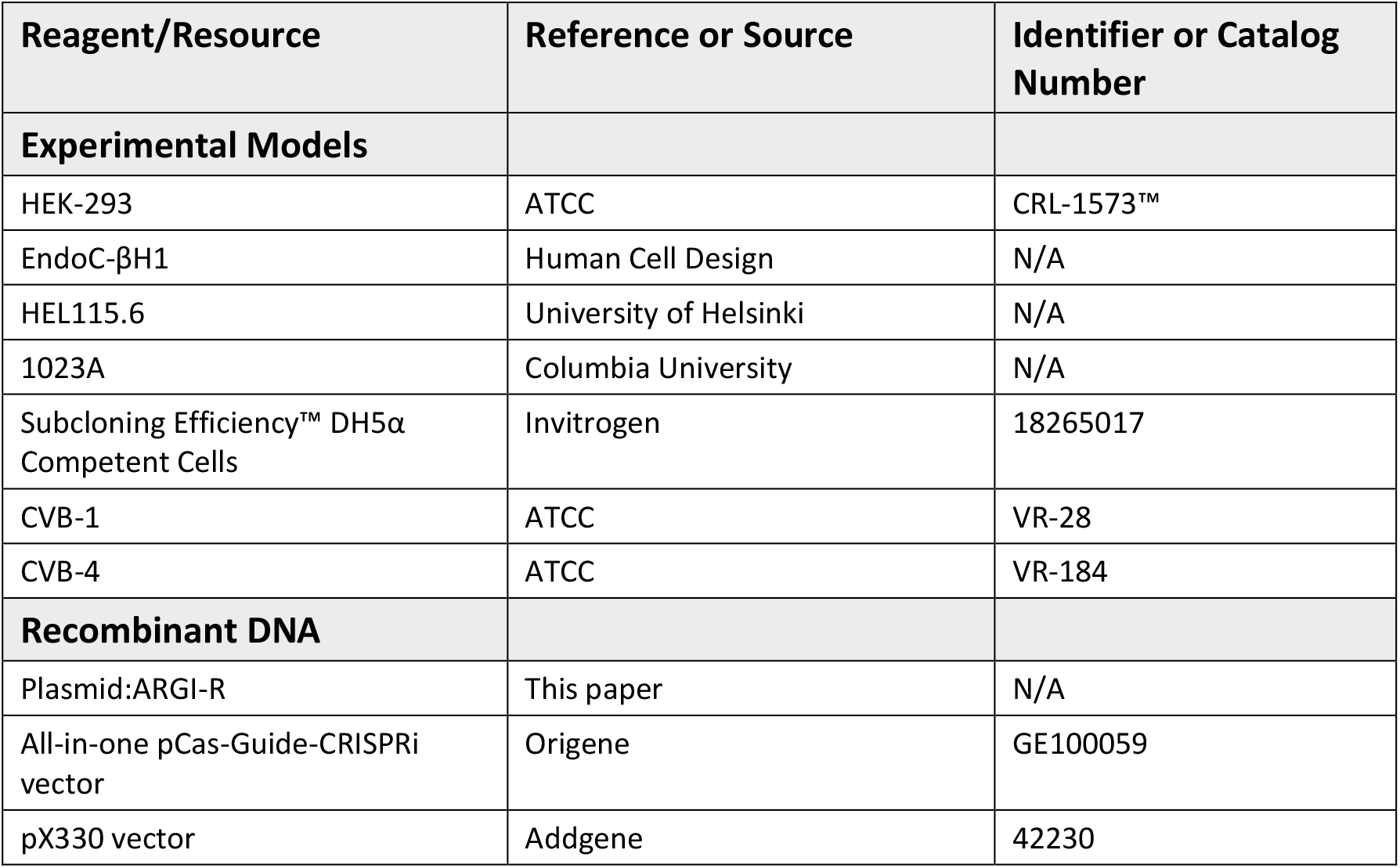
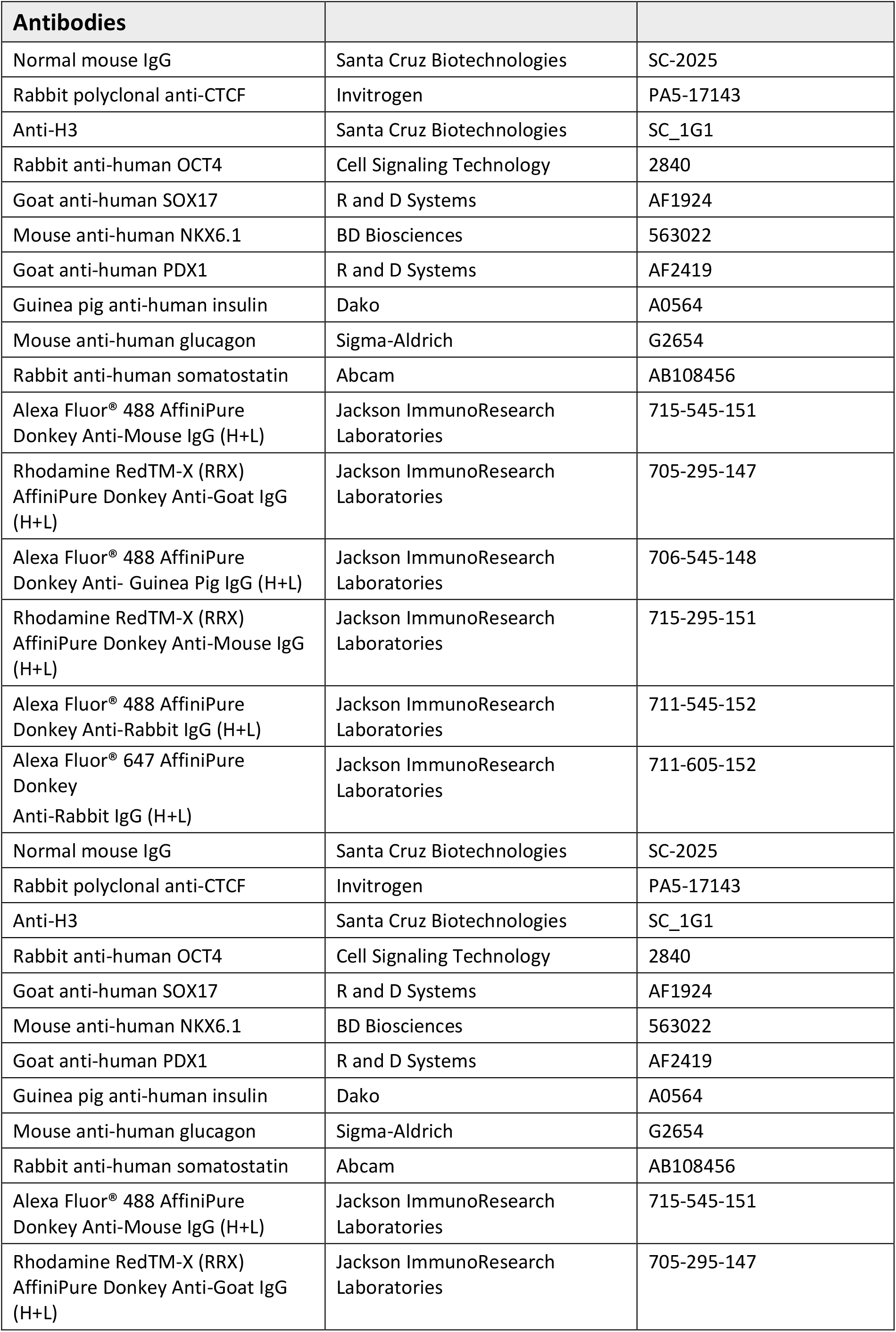
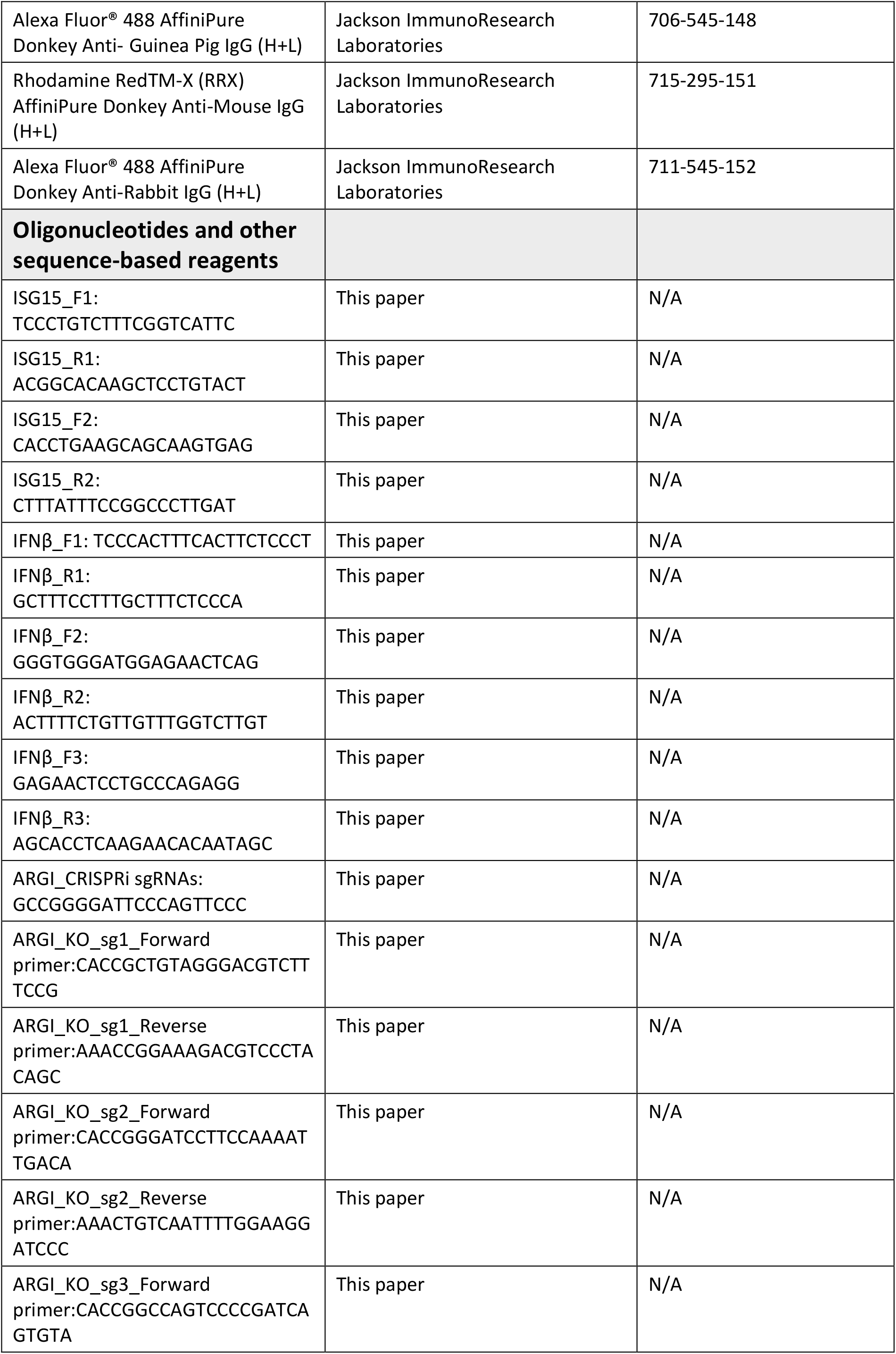
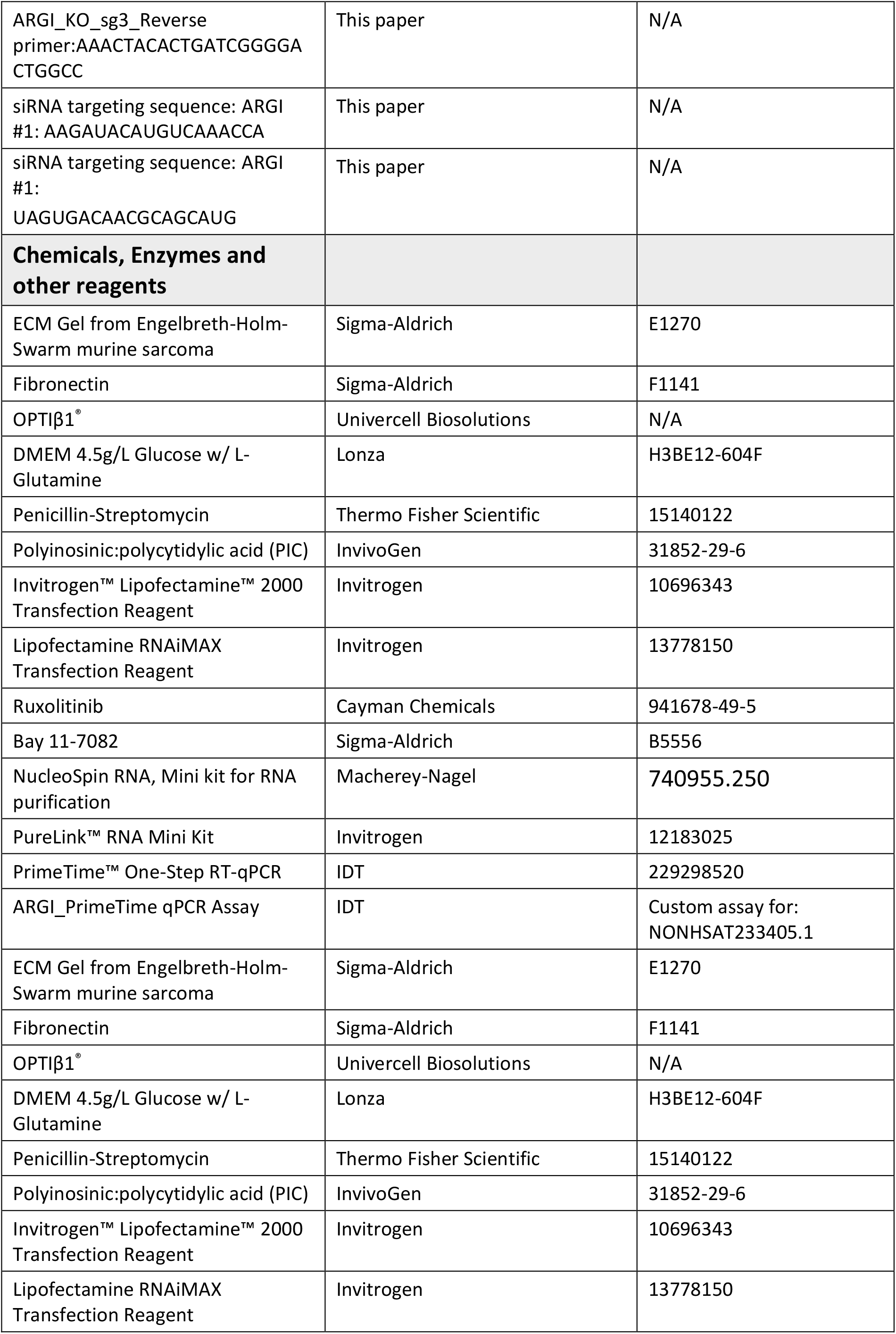
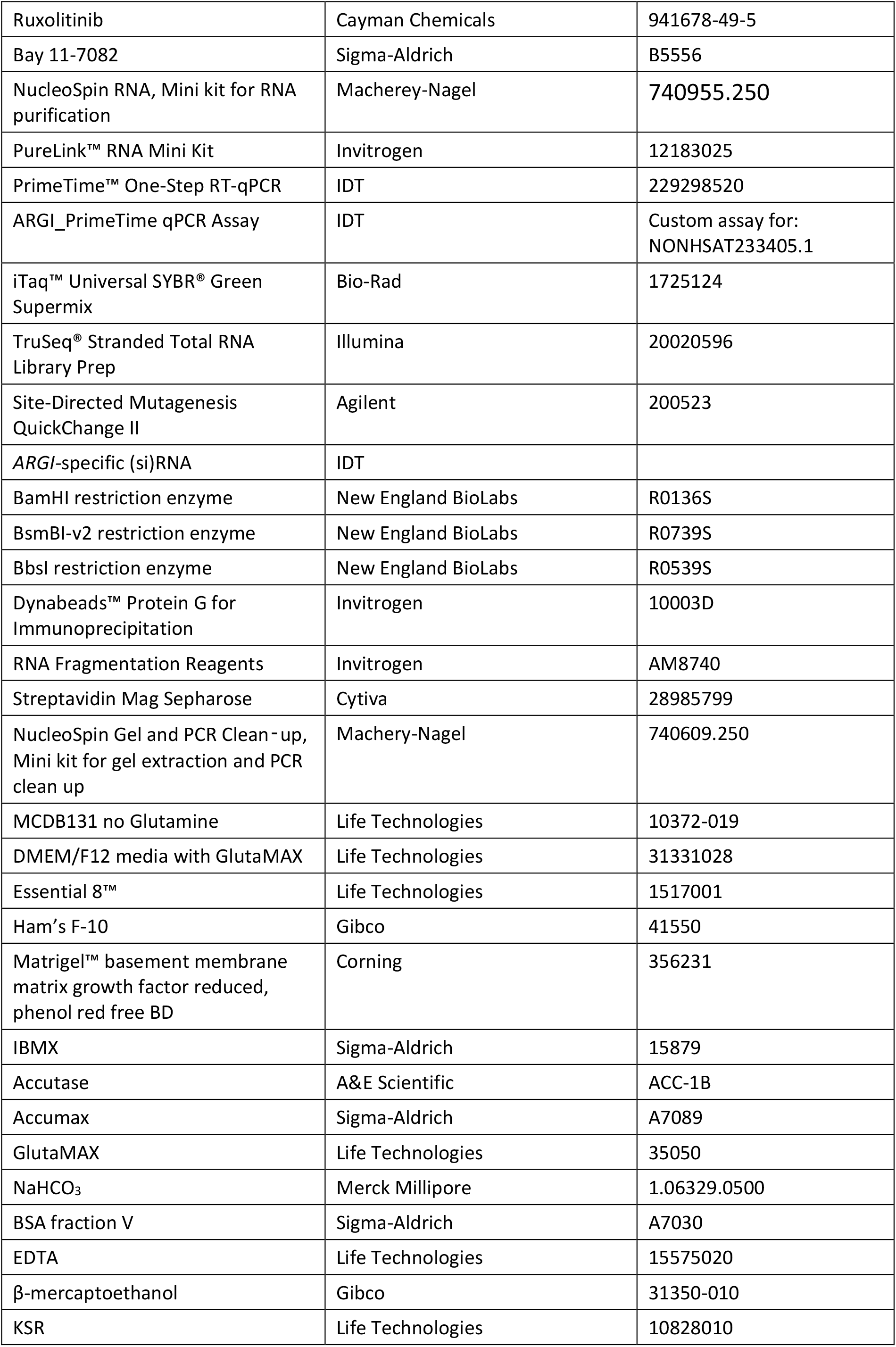
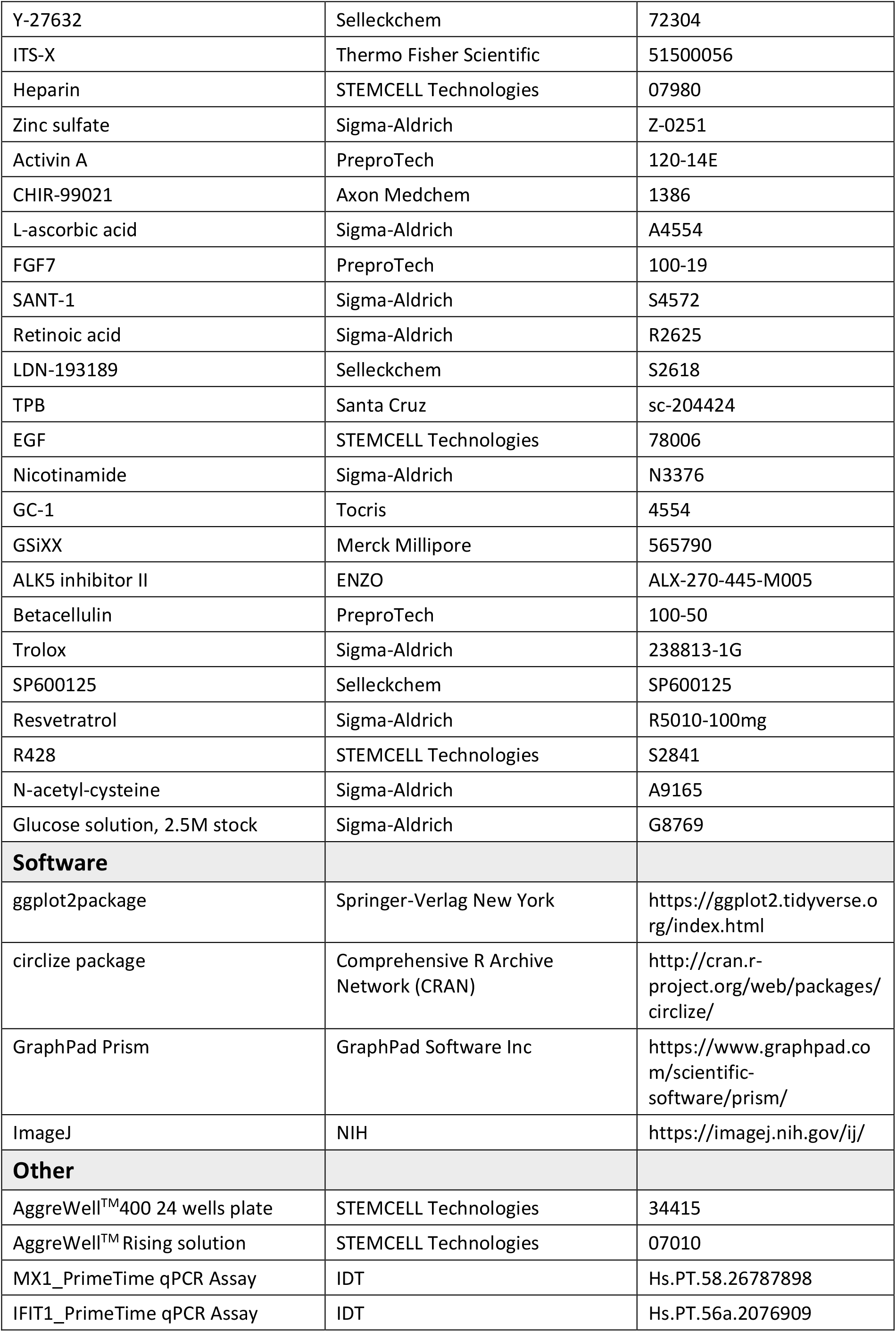
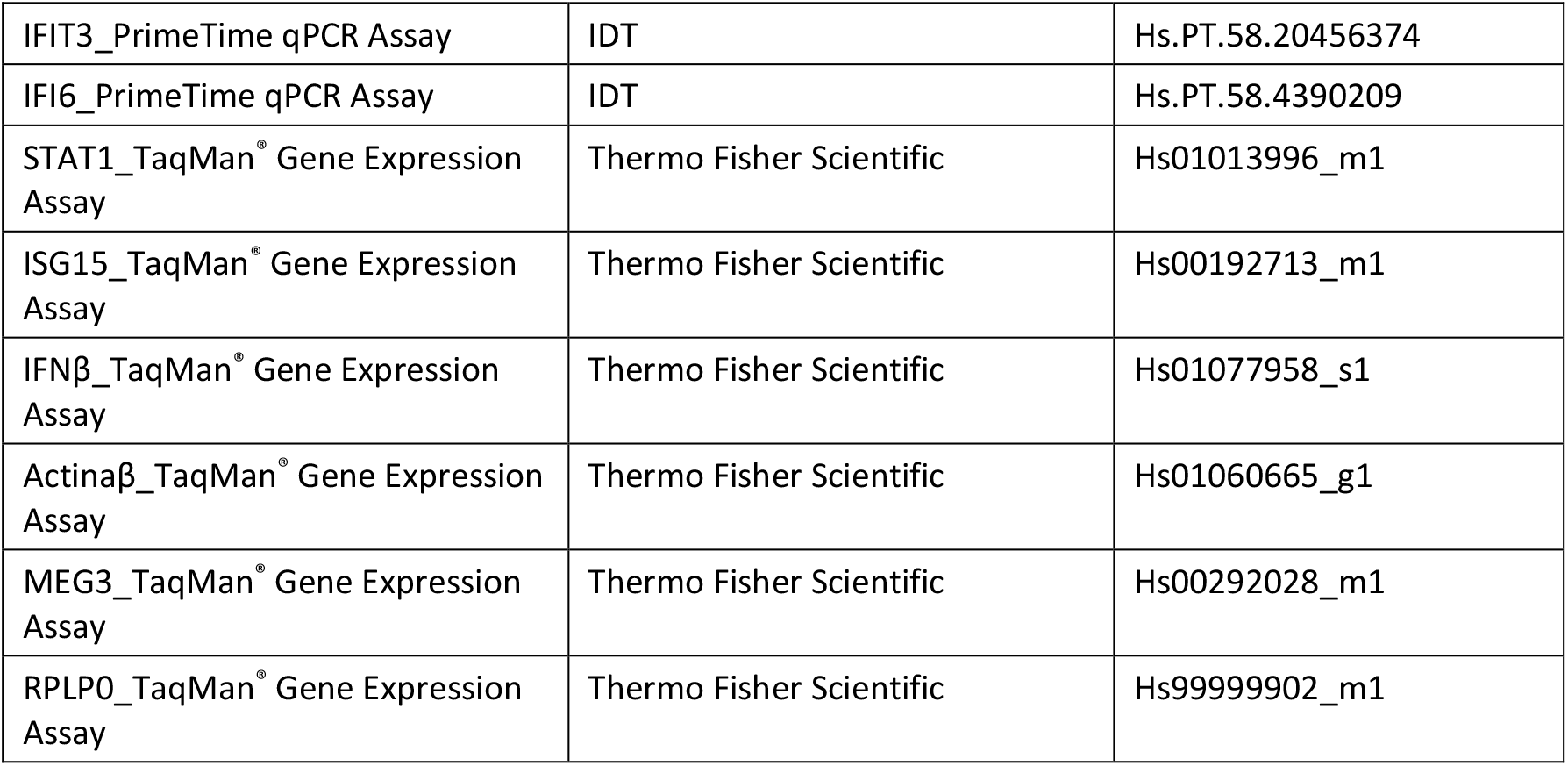

## Methods and protocols

### Human cell lines

The EndoC-βH1 human pancreatic β cell line was purchased from Human Cell Design (https://www.humancelldesign.com). Cells were cultured and split as previously described^60^. Briefly, EndoC-βH1 cells were seeded at an approximate density of 70.000-75.000 cells/cm^2^ on culture plates pre-coated with Matrigel-fibronectin (100mg/ml and 2 mg/mL, respectively, Sigma-Aldrich) at 37°C, 5% CO_2_ in complete OPTIβ1 medium (Human Cell Design). Cells were passed every 7 days. For transfection DMEM medium was used containing 2% FBS, 5.6 mmol/l glucose, 50 μmol/l 2-mercaptoethanol (Biorad, CA, USA), 10 mmol/l nicotinamide (Calbiochem, Darmstadt, Germany), 5.5 μg/ml transferrin, 6.7 ng/ml selenite (Sigma-Aldrich).

HEK293 cells (CRL-1573) were purchased from the American Type Culture Collection (ATCC; https://www.atcc.org). Cells were cultured in DMEM supplemented with 10% FBS and 100 units/ml penicillin and 100 μg/ml streptomycin (Lonza). The same medium without antibiotics was used for transfection.

### iPSC-derived human β cells

Two human iPSC lines (HEL115.6 and 1023A) derived from healthy donors were cultured and differentiated into β cells as described previously^61^ [HEL115.6 iPSCs were generated at the University of Helsinki; 1023A iPSCs were kindly provided by Dr DM Egli (Columbia University). The iPSCs had normal karyotype, stem cell colony morphology and expressed pluripotency markers^61^. iPSCs were maintained in E8 medium on Matrigel-coated plates (Corning) and seeded at 2.5–2.8 × 10^6^ cells per 3.5 cm well in E8 medium containing 5 μM ROCK inhibitor (STEMCELL Technologies) 24h prior to the 7-stage differentiation. When reaching pancreatic progenitor stage, cells were plated into 24 well microwell plates at a density of 750 cells/microwell (AggreWell, STEMCELL Technologies) with 10 μM ROCK inhibitor and 1μl/ml heparin (STEMCELL Technologies) to allow 3-dimensional formation of aggregates. Prior to infection, stage 7 aggregates were dispersed. Aggregates were incubated in 0.5 mM EDTA at room temperature for 4 min, exposed to Accumax (Sigma–Aldrich) for 8 min and then dispersed by gentle pipetting^62^. Knockout serum (Gibco) with 10 μM ROCK inhibitor was added to quench the dissociation process, and cells were seeded at 5 × 10^4^ cells per 6.4 mm well in stage 7 medium^61^.

### PIC transfection, viral infection and cell treatments

The synthetic viral dsRNA mimic PIC (HMW, InvivoGen) was used at a concentration of 1 μg/ml and transfected using Lipofectamine-2000 (Invitrogen) as previously described^3^.

For viral infection, iPSC-derived β cells were dispersed using Accumax (Sigma) as previously described^64^. Cells were plated at a cell density of 300,000/well in Matrigel-coated 24-well plates in stage 7 medium^61,64^ supplemented with 10 μM ROCK inhibitor. After overnight recovery, cells were infected with CVB1/Conn-5 (MOI=0.05) and CVB4/JVB (MOI=0.5) in Ham’s F-10 Nutrient Mixture (Gibco), supplemented with 2 mM GlutaMAX, 50 μM IBMX and 1% FBS. Two hours after infection medium was replaced by Ham’s F-10 Nutrient Mixture, 0.75% BSA, 2 mM GlutaMAX, 50 μM IBMX, 50 U/ml penicillin, 50 μg/ml streptomycin and cells were cultured for an additional 22h as previously described^31^.

The NFκB inhibitor Bay 11-7082 (Sigma-Aldrich) was used at a concentration of 10 μM.

### RNA extraction and quantitative PCR

RNA extraction was performed using the NucleoSpin RNA Kit (Macherey Nagel), PureLink RNA Mini kit (Invitrogen) or miRNeasy mini kit (Qiagen) and expression values were determined by qPCR using Taqman Gene Expression Assays (Thermo Scientific) specific for *ARGI, MX1, ISG15, IFI6, IFIT1, IFIT3, STAT1, MEG3, RPLP0* and *IFNβ* or by Sybr Green (Biorad) using specific primers for *ARGI, ISG15* promoter, *ISG15* enhancer and *IFNβ* promoter. The Taqman Gene Expression Assays and primers are listed in the Key resources table.

qPCR measurements were performed in duplicate in at least 3 independent samples and expression levels were analyzed using the 2^−ΔΔCt^ method.

### Cellular fractionation

For quantification of *ARGI* RNA levels in whole cell and nuclear extracts, nuclei were isolated using C1 lysis buffer (1.28 M sucrose, 40 mM Tris-HCl pH 7.5, 20 mM MgCl2, 4% Triton X-100). *ARGI, MEG3 or MALAT1* (as nuclear controls) and *RPLP0* (cytoplasmic control) levels were quantified by qPCR and compared to the total amount of those RNAs in the whole cell lysate.

### RNA-sequencing

RNA-sequencing of control and ARGI-overexpressing pancreatic β cells was performed in a NovaSeq 6000 sequencer. Sequencing libraries were prepared using “TruSeq Stranded Total RNA Human” kit (Illumina Inc., Cat.# RS-122-2201), following “TruSeq Stranded Total RNA Sample Prep-guide (Part # 15031048 Rev. E)”.

Reads were trimmed using Trimmomatic 0.39, with default settings. Then, reads were aligned by means of HISAT2 using as reference Human Genome assembly hg38, as it is available from the developer site. Stringtie was used to calculate transcripts and their abundances; and htseq-count to get the counts of each transcript.

Transcripts with more than 10 counts in at least 3 samples were kept. Then, counts were adjusted by means of RUVSeq package of R language, and *C1orf43, EMC7, PSMB2, PSMB4, RAB7A, REEP5, VPS29* genes were used as negative controls in the estimation of the factors of unwanted variation. edgeR package in R language was used to calculate differential expression between control and *ARGI* overexpression, using the upper quartile method for normalization and considering that samples were paired. Gene-set enrichment analyses were carried out by means of enrichR, and additional statistics analyses and graphs were made using R language base functions, ggplot2 package and cyclize package.

### Western blot analysis

EndoC-βH1 cells were washed with cold PBS and lysed in Laemmli buffer (62 mM Tris-HCl, 100 mmol/l dithiothreitol (DTT), 10% vol/vol glycerol, 2% wt/vol SDS, 0.2 mg/ml bromophenol blue, 5% vol/vol 2-mercaptoethanol). Proteins in the lysate were separated by SDS-PAGE. Following electrophoresis, proteins were transferred onto nitrocellulose membranes using a Transblot-Turbo Transfer System (Bio-Rad) and blocked in 5% wt/vol non-fatty milk diluted in TBST (20 mM Tris, 150 mM NaCl and 0.1% vol/vol Tween 20) at room temperature for 1h. The membranes were incubated overnight at 4°C with a primary antibody specific for CTCF (Cat #PA5-17143, Invitrogen) diluted 1:1000 in 5% wt/vol BSA or anti-GAPDH (SC-365062, Santa Cruz Biotechnologies) diluted 1:5000 in 5% wt/vol BSA. Immunoreactive bands were revealed using the Clarity Max Western ECL Substrate (Bio-Rad) after incubation with a horseradish peroxidase-conjugated anti-rabbit (1:1000 dilution in 5% wt/vol non-fatty milk) or anti-mouse (1:5,000 dilution in 5% wt/vol non-fatty milk) secondary antibody for 1h at room temperature. The immunoreactive bands were detected using a Bio-Rad Molecular Imager ChemiDoc XRS and quantified using ImageLab software (Bio-Rad).

### ARGI overexpression and silencing experiments

The overexpression vector for *ARGI* harboring the T1D risk allele (ARGI-R) was purchased from ProteoGenix (Schiltigheim, France). The overexpression vector for *ARGI* harboring the T1D protective allele (ARGI-P) was produced by site-directed mutagenesis using the Site-Directed Mutagenesis QuickChange II (Agilent). *ARGI*-overexpressing plasmids were transfected using Lipofectamine-2000 (Invitrogen) following the manufacturer’s instructions.

*ARGI* was silenced by transfection of a small interfering (si)RNA (see Key resources table) using Lipofectamine RNAi Max (Invitrogen) following the manufacturer’s instructions.

### CRISPRi and CRISPR-Cas9 experiments

For CRISPR-Cas9 experiments sgRNAs (listed in the Key resources table) were designed using the Zhang lab tool (currently Synthego) and cloned into the pX330 vector (Addgene) using BbsI restriction site (NEB). Plasmids harboring the sgRNAs were transfected into cells using Lipofectamine-2000. PIC was transfected as described above 3 days post sgRNA transfection.

For CRISPRi experiments a sgRNA (see Key resources table) was cloned into a CRISPRi vector harboring the dCas9 protein (Origene) using BamHI and BsmBI restriction sites. The plasmid harboring the sgRNA was transfected into cells using Lipofectamine-2000 and PIC was transfected as above 3 days post sgRNA transfection.

### RNA immunoprecipitation assay

For RNA immunoprecipitation experiments, EndoC-βH1 cells were kept untransfected or transfected with 1 μg/ml of PIC using Lipofectamine-2000 for 24h. Cells were lysed in RNA immunoprecipitation buffer (150 mM KCl, 25 mM Tris, 0.5 mM DTT, 0.5% NP-40, protease inhibitors), kept on ice for 15 minutes and homogenized using a syringe. Lysates were pre-cleared with Dynabeads G (ThermoFisher) for 1h in a wheel shaker at 4°C. Pre-cleared lysates were incubated with an anti-IgG antibody (negative control; Santa Cruz Biotechnologies) or anti-CTCF antibody (ThermoFisher) for 1h at room temperature in a wheel shaker. After, Dynabeads were added and the mix further incubated for 30 min in the wheel shaker. Supernatants were removed and beads washed three times with RNA immunoprecipitation buffer, three times with low salt buffer (50mM NaCl, 10mM Tris-HCl, 0.1% NP-40), three times with high salt buffer (500mM NaCl, 10mM Tris-HCl, 0.1% NP-40), and then resuspended in RNA extraction buffer.

The same procedure was used for immunoprecipitation of CTCF-bound *ARGI* in ARGI-R- or ARGI-P- and CTCF-overexpressing HEK293 cells.

### RNA antisense purification

RNA antisense purification was performed following a protocol from the Guttman lab^65^. In brief, probes were generated by *in vitro* transcribing biotinylated antisense *ARGI* followed by controlled RNA Fragmentation Reagent (Invitrogen). Probes against a non-related lncRNA of similar size were used as control. Extracts from EndoC-βH1 cells were crosslinked adding 2 mM disuccinimidyl glutarate for 45 min at room temperature and subsequently 3% formaldehyde for 10 min at 37°C. Crosslinked RNA-chromatin was fragmented by sonication, hybridized with control or *ARGI*-specific RNA probes, captured using Streptavidin Mag Sepharose (Cytiva), washed extensively and eluted. Retrieved RNA was purified using PureLink Micro RNA columns (Thermo Scientific) and fragmented DNA by NucleoSpin Gel and PCR Clean-up (Macherey-Nagel). Enrichment of *ARGI* in the regulatory regions of IFNβ and ISG15 was quantified by qPCR.

### Chromatin RNA immunoprecipitation

HEK293 cells were transfected with PIC, fixed with disuccinimidyl glutarate and formaldehyde; chromatin was sheared by sonication and immunoprecipitated using protein G magnetic Dynabeads and CTCF antibody. Immunoprecipitated material was eluted, reverse crosslinked and treated with Proteinase K. RNA was extracted using PureLink Micro RNA columns and DNA was extracted using the NucleoSpin Gel and PCR Clean-up. Immunoprecipitated RNA and DNA analyzed by qPCR using primers specific for *ARGI* and regulatory regions of IFNβ and ISG15.

### Quantification and statistical analysis

Experimental data are displayed as means±SEM. Comparisons between groups were performed by Student’s t test or one-way ANOVA followed by correction for multiple comparisons (as recommended by GraphPad Prism v8.0.1). p-values below 0.05 were considered statistically significant.

## Acknowledgments

This work was supported by grants from the Ministerio de Ciencia, Innovación y Universidades (PID2019-104475GA-I00), the European Foundation for the Study of Diabetes (EFSD) - EFSD/JDRF/Lilly Programme on Type 1 Diabetes Research to IS, the Fondation Franconphone pour la Recherche sur le Diabète (FFRD) to MC and MIE. MC acknowledges support by the Walloon Region SPW-EER (Win2Wal project BetaSource) and the Fonds National de la Recherche Scientifique (FRS-FNRS). IGM is supported by a Predoctoral Fellowship Grants from the UPV/EHU (Universidad del País Vasco/Euskal Herriko Unibertsitatea). AOG is supported by a post-doctoral grant from UPV/EHU (ESPDOC21/56, Universidad del País Vasco/Euskal Herriko Unibertsitatea), MNA is a FRIA F.R.S-FNRS fellow. TS is supported by a Marie Skłodowska-Curie Actions Fellowship from the European Union’s Horizon 2020 research and innovation programme under the Marie Skłodowska-Curie grant agreement No 801505

## Author contributions

I.S. contributed to the original idea, design the experiments, contributed to research data, and wrote the manuscript. I.G.M. contributed to research data and wrote the manuscript. K.G.E. and N.F.J. contributed to data analysis. L.M.M., M.N.A., A.O.G., T.S., M.C. A.O.B. and M.I.E. contributed to research data. M.C. edited the manuscript. All authors discussed the results, contributed to data interpretation, and approved the final manuscript.

