## Supplementary Figures for "The type 1 diabetes-associated lncRNA *ARGI* participates in virus-induced pancreatic β cell inflammation"

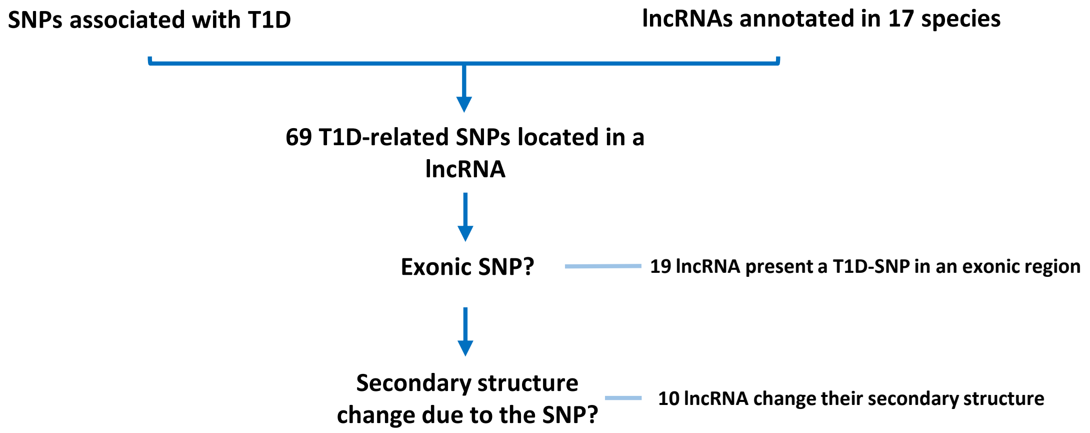


**Figure EV1. Schematic outline of the workflow used for T1D-associated lncRNA identification and selection.** Genomic positions of T1D-associated SNPs annotated in the NHGRI-EBI Catalog of Human Genome Association Studies (EMBL-EBI) were intersected with the genomic localization of all lncRNAs annotated in NONCODE version 6. Nineteen T1D-associated lncRNAs harboring an exonic SNP were analyzed in ViennaRNA Web Services to determine potential changes in their secondary structure. Ten lncRNAs were predicted to undergo secondary structure changes due to the T1D-associated SNP.

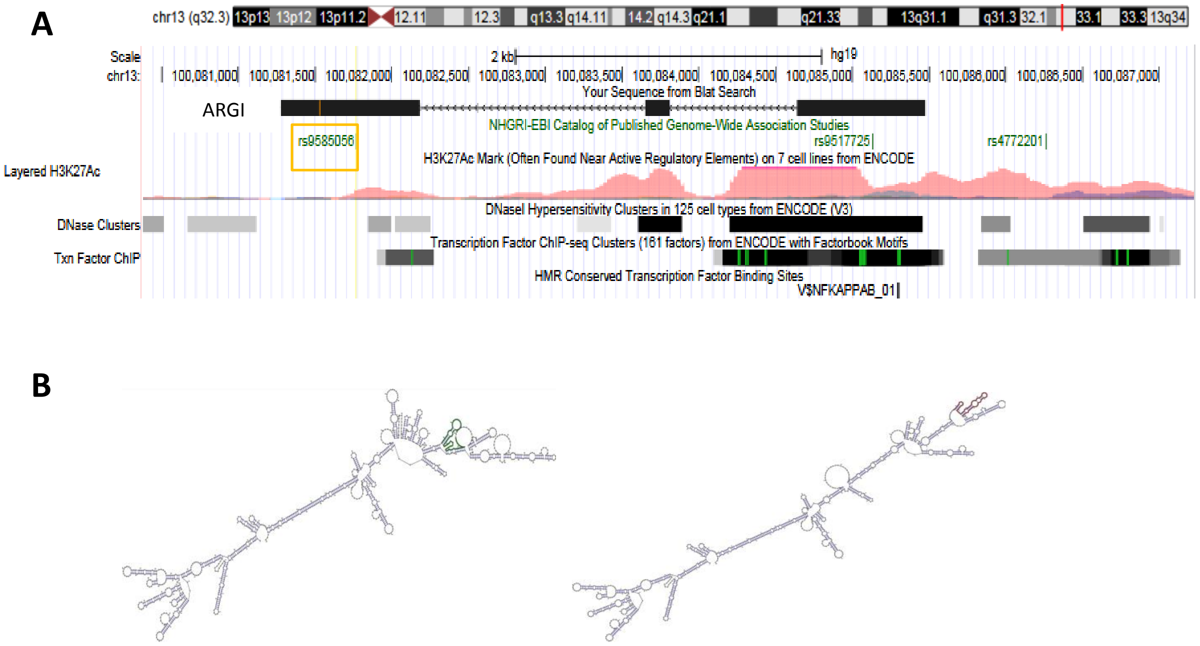


**Figure EV2. Genomic localization and secondary structure of *ARGI*.** (A) *ARGI* is an intergenic lncRNA located in human chromosome 13 (99,429,023-99,433,220; GRCh38/hg38). It has three exons and harbors one T1D-associated SNP (rs9585056; chr13: 99,429,262-99) in its third exon (orange box). Epigenetic marks (H3K27Ac and DNase clusters) and a conserved NFkB binding site have been identified close to the transcription start site of *ARGI*. (B) Secondary structure of *ARGI* predicted using the ViennaRNA Web Services. The image shows the secondary structure of *ARGI* harboring the T1D protective (rs9585056-A; green) or risk allele (rs9585056-G; red). There is a significant difference in the *ARGI* secondary structure prediction for the risk allele compared to the protective one (p=0.0536; the software considers significant whether p<0.2).

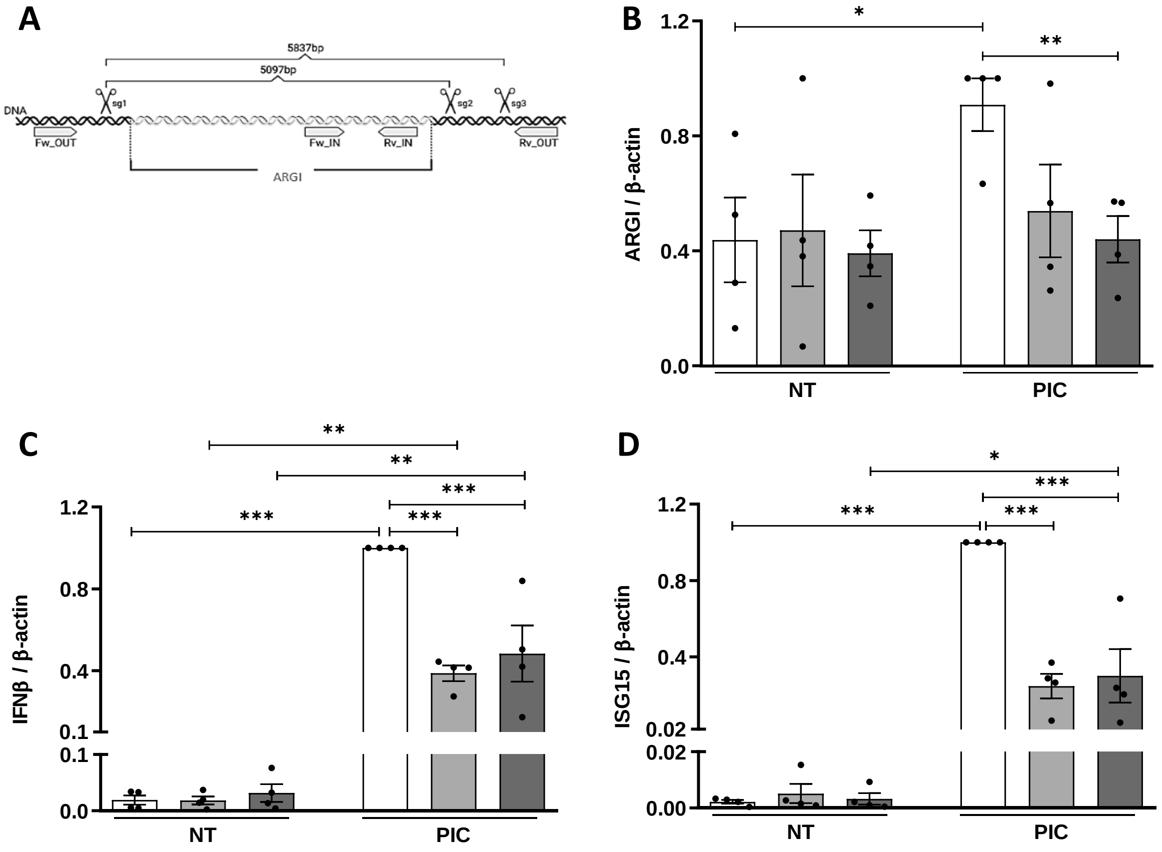


**Figure EV3. *ARGI* disruption using CRISPR-Cas9 reduces PIC-induced *IFN*b and *ISG15* expression.** (A) *ARGI* disruption was performed by generating a deletion of 5097 bp using single guide RNAs (sgRNAs) 1 and 2, or a deletion of 5837 bp using sgRNAs 1 and 3. The deletion was confirmed by PCR using a primer pair located inside the deleted region (for detection of unedited cells; wild type forward (Fw_IN) and wild type reverse (Rv_IN)) and a primer pair located outside the deleted region (for detection of edited cells; Fw_OUT and Rv_OUT). (B-D) EndoC-βH1 cells were transfected with an empty px330 vector (white bars) or with vectors harboring any of the two combinations of sgRNAs targeting *ARGI* (light and dark grey bars). After 36h, cells were left non-transfected (NT) or transfected with PIC (1 µg/ml) for 24h. Expression of *ARGI* (B), *IFN*b (C) and *ISG15* (D) was determined by qPCR and normalized to the reference gene β-actin. Results are means±SEM of 4 independent experiments; ***p < 0.001, **p < 0.01 and *p < 0.05 as indicated; ANOVA followed by Student's t test.

A

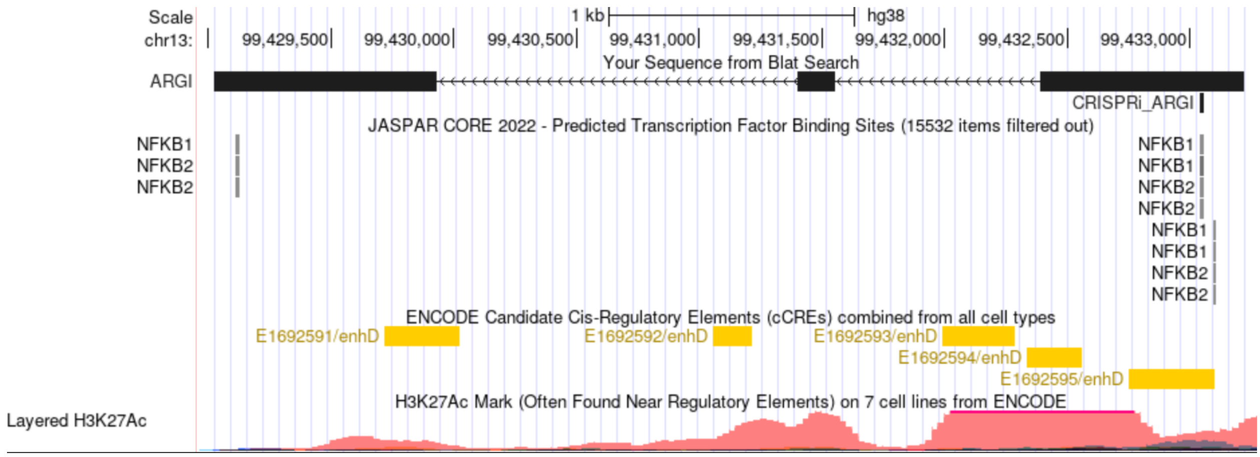


B

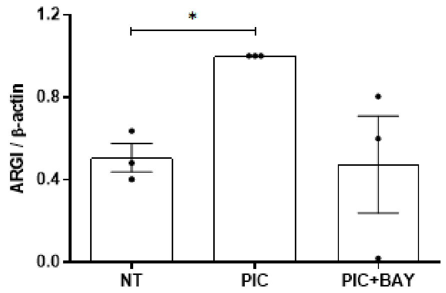


**Figure EV4. Inhibition of NFkB signaling counteracts PIC-induced *ARGI* upregulation in pancreatic β cells**. (A) Genomic location of the CRISPRi guide designed for the inhibition of *ARGI*. The guide is complementary to a conserved NFkB binding site located close to the *ARGI* transcription start site. (B) Human EndoC-bH1 cells were left untreated (NT), treated with intracellular PIC (1 µg/ml) for 24h (PIC) or treated with PIC and Bay 11-7082 (PIC+BAY). *ARGI* expression was determined by qPCR and normalized to the reference gene b-actin. Results are means±SEM of 3 independent experiments; *p < 0.05; Student's t test

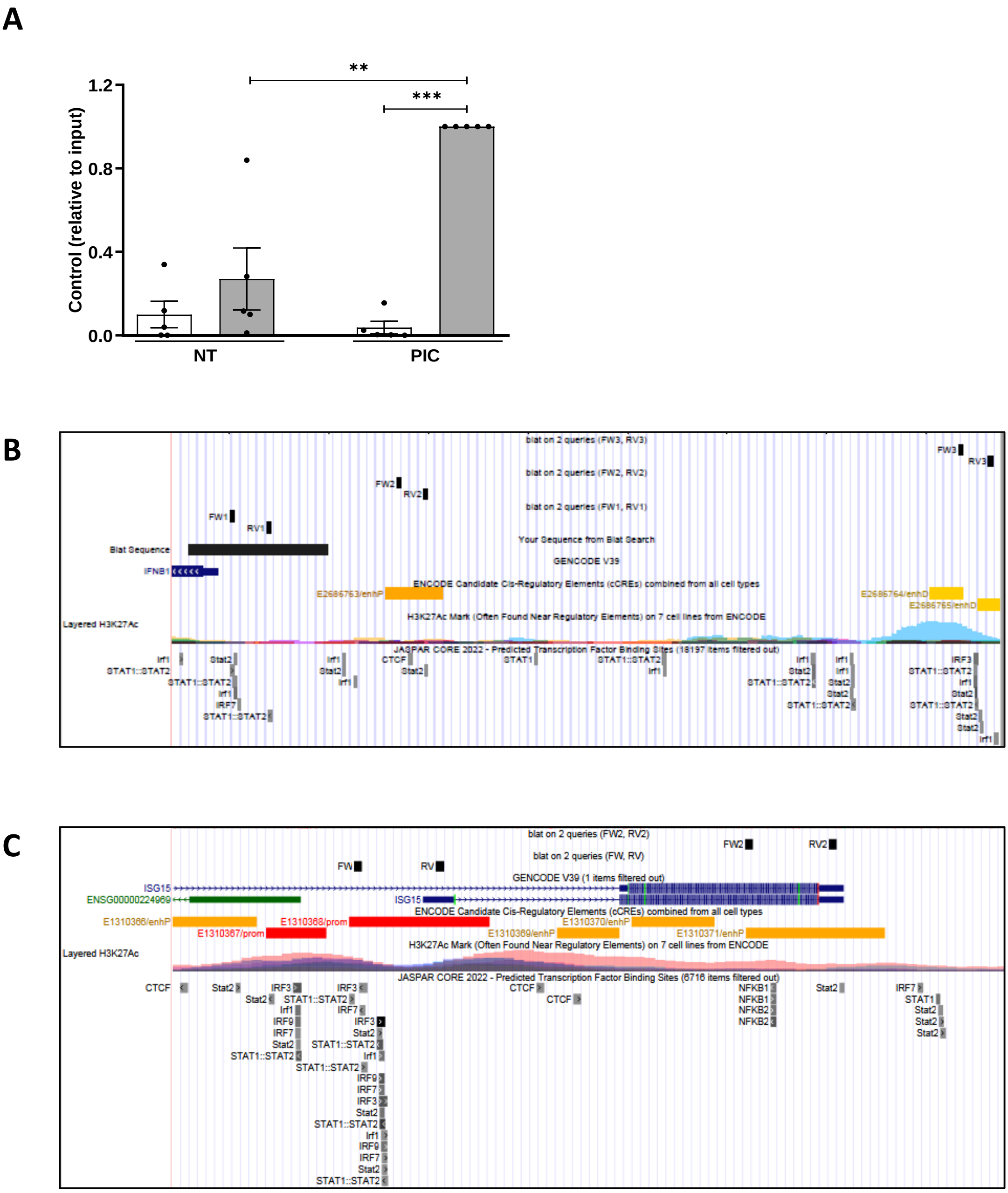


**Figure EV5. *ARGI* binds to the regulatory regions of *IFNb* and *ISG15* genes upon viral insult.** (A) RNA antisense purification of *ARGI* was performed in non-transfected (NT) or PIC-transfected EndoC-βH1 cells using probes targeting *ARGI* (white bars) or probes against a similar non-relevant lncRNA used as negative control (grey bars). Expression of the irrelevant control lncRNA was measured by qPCR. Results are presented as relative to input and are means±SEM of 5 independent experiments. ***$p < 0.001$ and **$p<0.01$ as indicated; ANOVA followed by Student's t test. (B-C) The location of the primer pairs used for the amplification of each region in the genomic map. (B) Primers FW1+RV1 were used to amplify *IFNb* promoter, and primers FW2+RV2 and FW3+RV3 primer pairs were used to amplify two distal enhancers of *IFNb* . (C) Primers FW+RV were used to amplify *ISG15* promoter and primers FW2+RV2 were used to amplify a distal enhancer of *ISG15*.
